# Proliferation genes repressed by TGF-β are downstream of Slug/Snail2 in normal bronchial epithelial progenitors and are deregulated in COPD

**DOI:** 10.1101/674127

**Authors:** Chamseddine Ben Brahim, Charlotte Courageux, Ariane Jolly, Bérengère Ouine, Aurélie Cartier, Pierre de la Grange, Leanne de Koning, Pascale Leroy

## Abstract

Slug/Snail2 belongs to the Epithelial-Mesenchymal Transition (EMT)-inducing transcription factors involved in development and diseases. Slug is expressed in adult stem/progenitor cells of several epithelia, making it unique among these transcription factors. To investigate Slug role in human bronchial epithelium progenitors, we studied primary bronchial basal/progenitor cells in an air-liquid interface culture system that allows regenerating a bronchial epithelium. To identify Slug downstream genes we knocked down Slug in basal/progenitor cells from normal subjects and subjects with COPD, a respiratory disease presenting anomalies in the bronchial epithelium and high levels of TGF-β in the lungs. We show that normal and COPD bronchial basal/progenitors, even when treated with TGF-β, express both epithelial and mesenchymal markers, and that the epithelial marker E-cadherin is not a target of Slug and, moreover, positively correlates with Slug. We reveal that Slug downstream genes responding to both differentiation and TGF-β are different in normal and COPD progenitors, with in particular a set of proliferation-related genes that are among the genes repressed downstream of Slug in normal but not COPD. In COPD progenitors at the onset of differentiation in presence of TGF-β, we show that there is positive correlations between the effect of differentiation and TGF-β on proliferation-related genes and on Slug protein, and that their expression levels are higher than in normal cells. As well, the expression of Smad3 and β-Catenin, two molecules from TGF-β signaling pathways, are higher in COPD progenitors, and our results indicate that proliferation-related genes and Slug protein are increased by different TGF-β-induced mechanisms.

**Graphical Abstract:** 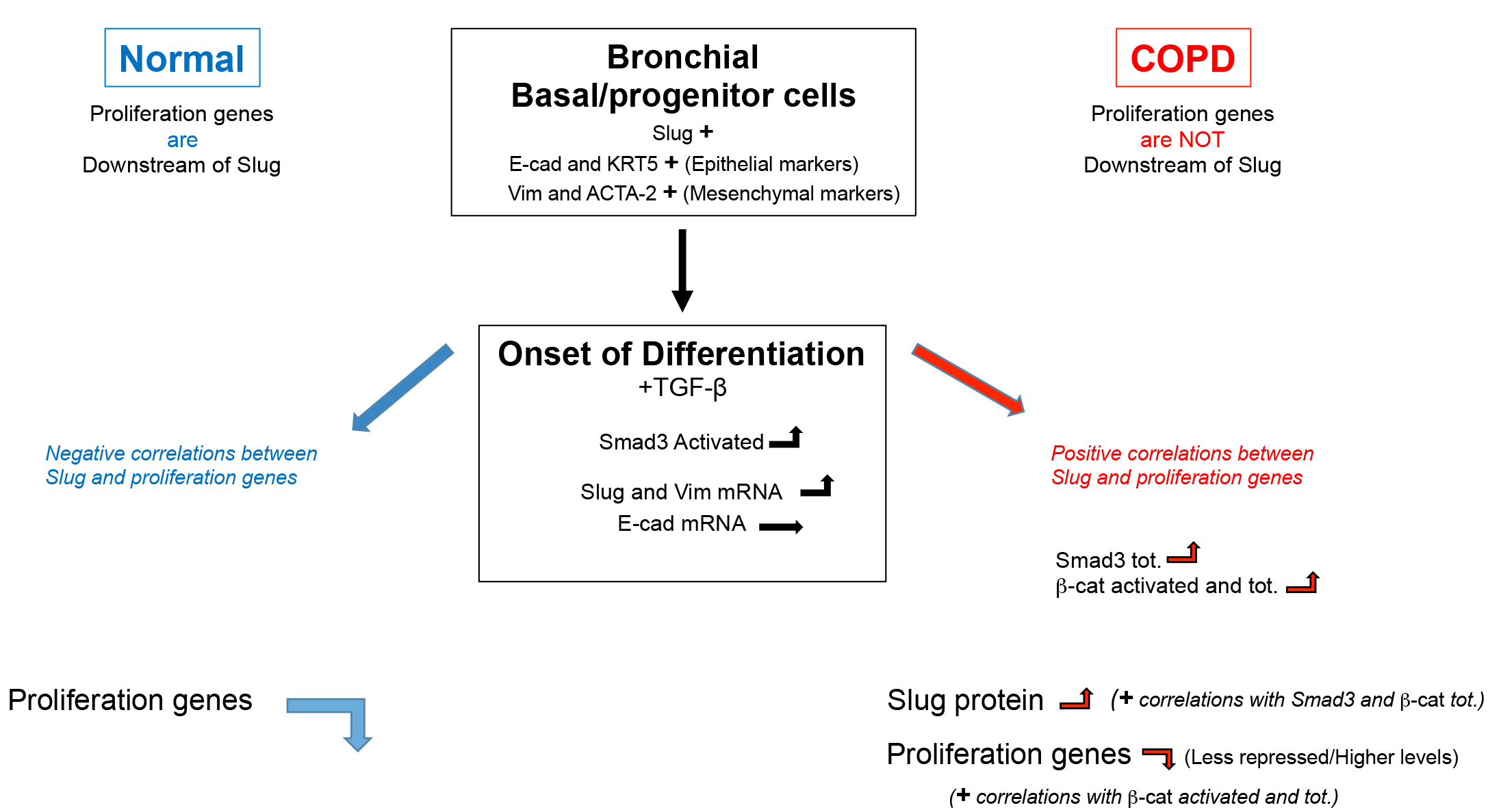

## Introduction

Slug (Snail2) belongs with Snail (Snail1) and Smuc (Snail3) to the Snail family of zinc finger transcription factors and has been studied essentially for its function as an inducer of Epithelial-Mesenchymal Transition (EMT) [1]. Like other EMT-inducing transcription factors it is expressed in a wide variety of tissues and cell types, during embryogenesis and organ formation\, and it is overexpressed and involved in EMT-related processes in numerous carcinoma. However, in contrast to other EMT-inducing transcription factors, Slug is expressed in adult stem/progenitor cells in normal epitheliums. Recently, it has been shown to control stem/progenitor cell growth during mammary gland morphogenesis [2] and to repress differentiation in epidermal progenitor cells [3]. In the lung, Slug is expressed in spheroids of stem/progenitor cells isolated from human adult airway epithelia [4] and, by microarray analysis, it was found enriched in both mouse and human airway basal cells [5-6].

Basal cells are the adult stem/progenitor cells of the airway epithelium: they can self-renew and/or differentiate to repair the epithelium after injury [7]. These cells can be isolated from bronchial tissue and, when grown in an air-liquid interface (ALI) culture system, reconstitute the pseudostratified bronchial epithelium [8]. In this system, basal cells keep the memory of the characteristics of their source tissue up to passage 3 to 4; e.g. when isolated from patients with Chronic Obstructive Pulmonary Disease (COPD), they reconstitute an epithelium with anomalies characteristic of COPD [9-10].

COPD is a respiratory disease characterized by a progressive and irreversible loss of respiratory capacity, with subjects at stage GOLD (Global initiative for chronic Obstructive Lung Disease) 1/A having mild symptoms and at GOLD 4/D having severe symptoms. One of the characteristics of COPD subjects is a remodeling of the airways with, in particular, many anomalies of the epithelium lining the airways such as basal cells hyperplasia, goblet cell metaplasia or squamous metaplasia [11-12]. Cigarette smoke is the main cause of COPD with 80-90% of patient developing COPD being smokers. Several studies have shown that cells exposed to cigarette smoke keep the memory of the exposure and establish a “field of injury” all along the airway epithelium [13]. The hypothesis is that the anomalies of the epithelium would result from an imbalance in the basal/progenitor cell fate, with this imbalance progressively worsening with the disease stage and leading to an increase of the extent of the anomalies. These deregulations are thought to be irreversible, since the anomalies are maintained even after subjects quit smoking [14]. Noticeably, COPD subjects have an increased risk of lung cancer compared to non-COPD smokers, in particular of squamous carcinoma in the proximal bronchus [15-16].

Our goal was to better understand Slug function in bronchial epithelium basal/progenitor cells. We hypothesized that Slug is involved in basal/progenitor cell fate i.e. self-renewal and differentiation, and that its deregulation therefore leads to anomalies in the bronchial epithelium. The ALI cell culture system allows regenerating, from primary basal cells, a bronchial epithelium with characteristics close to the in vivo source tissue, providing a good model to characterize the genetic program that takes place during differentiation.

COPD subjects present many anomalies in their bronchial epithelium that can be reproduced in the ALI cell culture system [10]. Moreover, features of EMT have been reported in COPD airways [9,17] and Transforming Growth Factor (TGF)-β is found at higher levels in COPD lung tissues [18]. TGF-β is known to regulate Slug expression [19], to be, in certain conditions, an EMT-inducing factor [20], and to play a role in stem/progenitor cell fate [21]. Hence, we studied basal/progenitor cells from COPD subjects and compared them with basal/progenitor cells from normal subjects in ALI cell culture system in conditions with or without TGF-β. We determined Slug expression and, since Slug is a transcription factor, we identified its downstream genes in bronchial epithelial progenitor cells, comparing cells from normal and COPD subjects at the onset of differentiation in presence or absence of TGF-β.

## Materials and methods

### Study subjects and cell isolation

Human lung tissues, non-COPD (non-smokers (n=6) and smokers (n=6)) and COPD (smokers (n=6) and Ex-smokers (n=6)) (see Appendix S1 for characteristics; COPD patients were diagnosed according to the GOLD guidelines) were obtained from patients undergoing lung lobectomy for peripheral lung carcinoma for removal of a primary lung tumor. Lung tissues used in this study were dissected as far away as possible from the tumor. Primary human bronchial epithelial cells were isolated from a piece of large bronchus according to standard protocol.

### Cell culture and ALI differentiation system

Primary human bronchial epithelial cells (Passage 1 or 2) were expanded on flasks coated with collagen I (BD Biocoat) in bronchial epithelial growth medium (BEGM), composed of bronchial epithelial basal medium (BEBM) supplemented with the SingleQuots kit (Lonza) and incubated at 37°C in 5% de CO2. Cells were expanded and differentiated in ALI (Air Liquid Interface condition) according to Fulcher et al [8] with the following modifications: Cells ≤90% confluent were plated in BEGM at a density of 1.5 × 10^5^ / cm2 on Transwell cell culture inserts with a polyethylene (PET) membrane and 0.4 μm pores (Corning Costar) coated with Collagen IV (Sigma). The next day, medium was replaced with ALI medium. Undifferentiated cells correspond to cells submerged and at day 3 post-plating. Cells at the onset of differentiation correspond to cells put in ALI condition (removal of the medium at the top of the culture) at day 3 post-plating and further cultured for 6 days. TGF-β1 (Peprotech) was added at a concentration of 1 ng/ml 2 days before the switch to ALI condition.

### shRNA lentivirus transduction

Protocol for transduction was obtained from D. Bryant [22] and adapted as following: Cells were plated at a density of 10^5^/cm2 in BEGM on Transwell cell culture inserts with a PET membrane and 0.4μm pores (Corning Costar) coated with Collagen IV (Sigma), and allow to adhere 3-4 h at 37°C, 5% CO2. shRNA lentiviral transduction particles from Mission RNAi TRC2 corresponding to SNAI2 specific or control non-targeting sequences (see Appendix S2 for sequence) were added on top of the cells at a MOI of 10. Inserts were swirled and placed O.N. at 37°C, 5% CO2. Medium was diluted by half with BEGM and insert placed back at 37°C, 5% CO2 for 1 day. Medium was replaced with ALI medium. At day 4 post-transduction, cells were changed to ALI conditions. Cells were lysed for RNA or protein analysis at day 6 post-transduction.

### Immunofluorescence staining of cell layers

All steps were done on a rocking table. Cells on inserts were rinsed with phosphate buffer saline (PBS)+ 1x, fixed with cold 4% paraformaldehyde in PBS+ for 10 min at 4°C, then permeabilized with 0.2% Triton-X100 in PBS for 10 min. For staining, blocking, incubations and washing were done with PFS (0.7% fish gelatin (Sigma) and 0.025% saponin in PBS 1x). Cells were first blocked with PFS for 30 min at room temperature, then incubated with primary antibodies at 4°C overnight. After extensive wash with PFS, cells were incubated at R.T. for 1 h with the corresponding Alexa Fluor conjugated secondary antibody at a dilution of 1:1000, then washed extensively. Cells were quickly rinsed twice with PBS 1x, adding Hoescht 33342 at 1 μg/ml in the first wash. Filters with the cells were cut out and mounted between a glass slide and a coverslip using ProLong gold (Invitrogen). Slides were left a minimum of 48 h at room temperature before observation under the microscope. Slides were frozen at - 20°C for long term storage. (see Appendix S3 for antibody references).

### RNA extraction

Cells were rinsed with phosphate buffer saline (PBS) before to be homogenized in RNA lysis buffer (NucleoSpin RNA Kit, Macherey-Nagel), supplemented with 1% β-Mercaptoethanol. Lysates were vortexed and proceeded immediately or snapped Frozen and stored at -80°C. Total RNA was extracted on column using the NucleoSpin RNA Kit (Macherey-Nagel). DNase I treatment was done directly on the column. RNA concentration was determined using a NanoDrop.

### cDNA synthesis

For each sample, 0.3 μg of total RNA was reverse transcribed as previously described [23] with the following modifications: Briefly, total RNA was annealed with 0.1 mg/ml oligo(dT)15 primer (Promega) and cDNA synthesis was performed with M-MLV Reverse Transcriptase (Promega) for 1 h at 42°C. A control without reverse transcriptase was done for each series of cDNAs.

### Quantitative Real-Time Polymerase Chain Reaction (qPCR)

Quantitative PCR was done on the QuantStudio 6 Flex with the QuantStudio Real-Time PCR software using SYBR green PCR master mix (Applied BioSystems). Primers (see Appendix S2 for primer sequences) were designed to be used with an annealing temperature of 60°C and an elongation time of 1 min. For a given gene target, cDNA volume used was chosen to be in a Ct range from 20 to 30, using 0.125 μM each forward and reverse primer. Glyceraldehyde-3-Phosphate Dehydrogenase (GAPDH) was used to normalize for cDNA amounts between samples.

### Protein extraction

Cells were rinsed with cold Tris 20 mM pH 7.5 and lysed directly on inserts with RIPA buffer (Tris pH 7.5, 50 mM; NaCl, 150 mM, NP-40 (leg10) or Triton X-100, 1%: Sodium deoxycholate, 1%; SDS 0.1%). Cells were scrapped and lysates were placed on a wheel for 30 min at 4°C. DNA was sheared with a needle and lysates spun down at 14K for 10 min at 4°C. Protein concentration in the lysates was determined using BCA protein assay (Pierce). Lysates were proceed immediately or snapped frozen and stored at -80°C.

### Reverse Phase Protein Array (RPPA)

RPPA was done by the RPPA platform at Curie Institute on samples prepared in RIPA buffer (as described in protein extraction) and stored at -80°C. RPPA was done as following: Protein concentration was determined (Pierce BCA reducing agent compatible kit, ref 23252). Samples were printed onto nitrocellulose covered slides (Supernova, Grace Biolabs) using a dedicated arrayer (2470 arrayer, Aushon Biosystems). Five serial dilutions, ranging from 1000 to 62.5 µg/ml, and four technical replicates per dilution were printed for each sample. Arrays were labeled with 32 specific antibodies, of which 8 were described in this study (see S3 Appendix for antibody references) or without primary antibody (negative control), using an Autostainer Plus (Dako). Briefly, slides were incubated with avidin, biotin and peroxydase blocking reagents (Dako) before saturation with TBS containing 0.1% Tween-20 and 5% BSA (TBST-BSA). Slides were then probed overnight at 4°C with primary antibodies diluted in TBST-BSA. After washes with TBST, arrays were probed with horseradish peroxidase-coupled secondary antibodies (Jackson ImmunoResearch Laboratories, Newmarket, UK) diluted in TBST-BSA for 1 h at RT. To amplify the signal, slides were incubated with Bio-Rad Amplification Reagent for 15 min at RT. The arrays were washed with TBST, probed with IRDye 800CW Streptavidin (LiCOR) diluted in TBST-BSA for 1 h at RT and washed again in TBST. For staining of total protein, arrays were incubated 30 min Super G blocking buffer (Grace Biolabs), rinsed in water, incubated 5 min in 0,000005% Fast green FCF (Sigma) and rinsed again in water. The processed slides were dried by centrifugation and scanned using a Innoscan 710-AL microarray scanner (Innopsys). Spot intensity was determined with MicroVigene software (VigeneTech Inc). All primary antibodies used in RPPA have been previously tested by Western Blotting to assess their specificity for the protein of interest. Raw data were normalized using Normacurve [24], which normalizes for fluorescent background per spot and a total protein stain.

### Gene expression profiling

Microarray analysis was performed on biological triplicate samples. Total RNA were amplified and labeled before hybridization onto Affymetrix human Gene 2.1 ST GeneChip according the manufacturer, by the Genomics platform at Curie Institute, Paris [25]. Array datasets were controlled using Expression console (Affymetrix) and further analyses and visualization were made using EASANA (GenoSplice, www.genosplice.com), which is based on the FAST DB annotations [26-27] Gene Array data were normalized using quantile normalization. Background corrections were made with antigenomic probes and probes were selected as described previously [28]. Only probes targeting exons annotated from FAST DB transcripts were selected to focus on well-annotated genes whose mRNA sequences are in public databases [26-27]. Bad-quality selected probes (e.g., probes labeled by Affymetrix as ‘cross-hybridizing’) and probes whose intensity signal was too low compared to antigenomic background probes with the same GC content were removed from the analysis. Only probes with a DABG P value ≤ 0.05 in at least half of the arrays were considered for statistical analysis [28]. Only genes expressed in at least one compared condition were analyzed. To be considered expressed, the DABG P value had to be ≤ 0.05 for at least half of the gene probes. We performed an unpaired Student’s t-test to compare gene intensities in the different biological replicates. Genes were considered significantly regulated when fold-change was ≥ 1.2 and P value ≤ 0.05. Significant KEGG pathways [29], REACTOME pathways [30] and GO terms were retrieved using DAVID [31] from union of results of all, up- and down-regulated genes separately. Data set GEO ID numbers are GSE122957 and GSE123129.

### Image capture and analysis

Images of cell layers stained by immunofluorescence were captured using a SP8 Leica confocal microscope equipped with a X 40 objective. The following excitation sources where used: a diode 405-nm, an argon laser 488-nm, a diode 561-nm, a diode 633-nm. Detectors were PMT and HyD. Digital images were analyzed using ImageJ software.

### Statistical Analysis

Biological replicates were n ≥ 11 and data generated by at least 2 independent experiments. For RNA or protein levels, mean is ±SD and statistical analysis was carried out by parametric paired or unpaired two-sided t-test as appropriate. For fold-change, mean is ±SEM and statistical analysis was carried out by a one sample two-sided t-test. Correlations were computed as Pearson correlation coefficients and P value determined by two-sided test. Significance was accepted when P value < 0.05.

## Results

### Slug/Snail2 is the only EMT-inducing transcription factor highly expressed in basal cells and it co-expresses with epithelial and mesenchymal markers

Slug/Snai2 gene has been found by microarray analysis to be highly expressed in basal cells of the human airway epithelium [5-6]. To determine the role of Slug in these stem/progenitor cells, we used primary basal cells isolated from human bronchial epithelium and first characterized these cells for the expression of Slug.

We confirmed by immunocytochemistry that primary basal cells grown at confluence and maintained undifferentiated are all progenitor cells as shown by the expression of the marker p63, and that they all express Slug, which co-localizes with p63 in their nuclei (Fig. 1a). Similar expression profiles were obtained with cells from COPD subjects (Fig. S1). Slug is an EMT-inducing transcription factor and we also determined the expression of other transcription factors with this property. While Slug is highly expressed, the expression of Snail1 and Zeb1 is close to background and Twist1 does not exceed 10% of Slug levels, and this in both normal and COPD cells (Fig. 1b). To determine the epithelial status of these progenitor cells, we studied genes coding for junction proteins and found that the epithelial marker E-cadherin (E-cad/CDH1) is expressed while N-cadherin (N-cad/CDH2), an EMT-related mesenchymal marker, is not (Fig. 1c). We also detected high levels of expression of the gene coding for the epithelial cytoskeletal protein cytokeratin 5 (KRT5) as well as of ACTA2, the gene coding for α-smooth muscle actin (a-sma) and vimentin (Vim), two mesenchymal cytoskeletal proteins (Fig. S2a). In conclusion, both normal and COPD progenitor cells express Slug as well as a combination of epithelial (E-cad, KRT5) and mesenchymal (Vim, ACTA-2) markers.

**Fig. 1.**
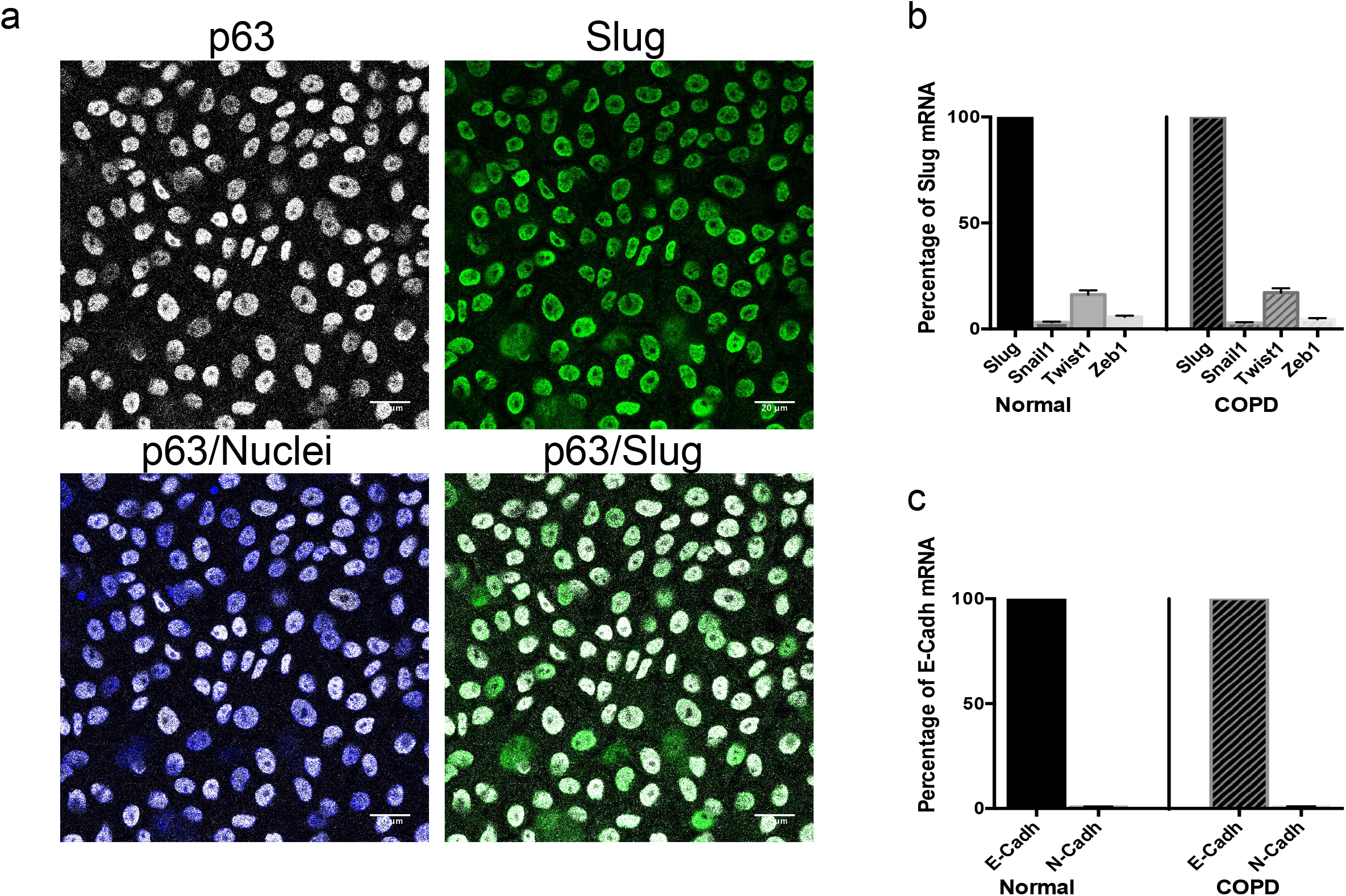
Bronchial epithelial progenitors express Slug in their nuclei and co-express epithelial and mesenchymal markers. Primary bronchial epithelial basal cells were grown on filters and analyzed undifferentiated at confluence, either by fluorescent immunocytochemistry (**a**) or by RT-qPCR (**b, c**) (**a**) Normal cells were fixed and labeled simultaneously with progenitor cell marker p63 (white) and Slug (green) antibodies and with Hoechst as a marker of nuclei (blue). Bars are 20 µm. (**b, c**) RNA were extracted from normal and COPD cells and analyzed by RT-qPCR. GAPDH was used to normalize cDNA amounts between samples and results were calculated as a ratio on GAPDH. Data are for n ≥ 11 and experiments were done at least in duplicate. (**b**) Expression of the EMT-inducing transcription factors Snai1, Twist1 and Zeb1 shown as a percentage of Slug mRNA, with mean ±SEM (**c**) Expression of the mesenchymal marker N-cad/CDH2 shown as a percentage of the epithelial marker E-cad/CDH1 mRNA, with mean ±SEM

### Slug/Snail2 and E-cadherin co-express in basal cells in absence or presence of TGF-β

Next, we sought to determine if the expression of Slug and the EMT-related markers is modified in cells in which differentiation is induced. For this, we used the ALI cell culture system that allows basal/progenitor cells to reconstitute a characteristic bronchial pseudostratified epithelium. In the ALI system, complete differentiation takes 3 to 4 weeks but we were interested in the onset of the genetic differentiation program and studied timepoints at day 6 of culture. TGF-β can play a role in stem/progenitor cell fate [21]; it also regulates Slug expression, is a potential EMT-inducing factor and is expressed at higher levels in COPD airways compared to normal [18-19]. We thus compared the expression of Slug and the EMT-related markers in absence or presence of TGF-β to determine if progenitor cells from normal and COPD responded differently to TGF-β. We used low concentration of TGF-β (1ng/ml) to be in conditions similar to the physiological range.

In both normal and COPD cells, TGF-β significantly increases Slug mRNA expression while its effect on the expression of E-cad/CDH1 is not statistically significant (Fig. 2a, b). Slug remains the most highly expressed EMT-inducing transcription factor and E-cad/CDH1 has still a much higher expression than the mesenchymal marker N-cad/CDH2 (Fig. 2c, d). Among the cytoskeletal markers, Vim is highly upregulated by TGF-β, while KRT5 and ACTA2 levels do not change significantly (Fig. S2b).

**Fig. 2.**
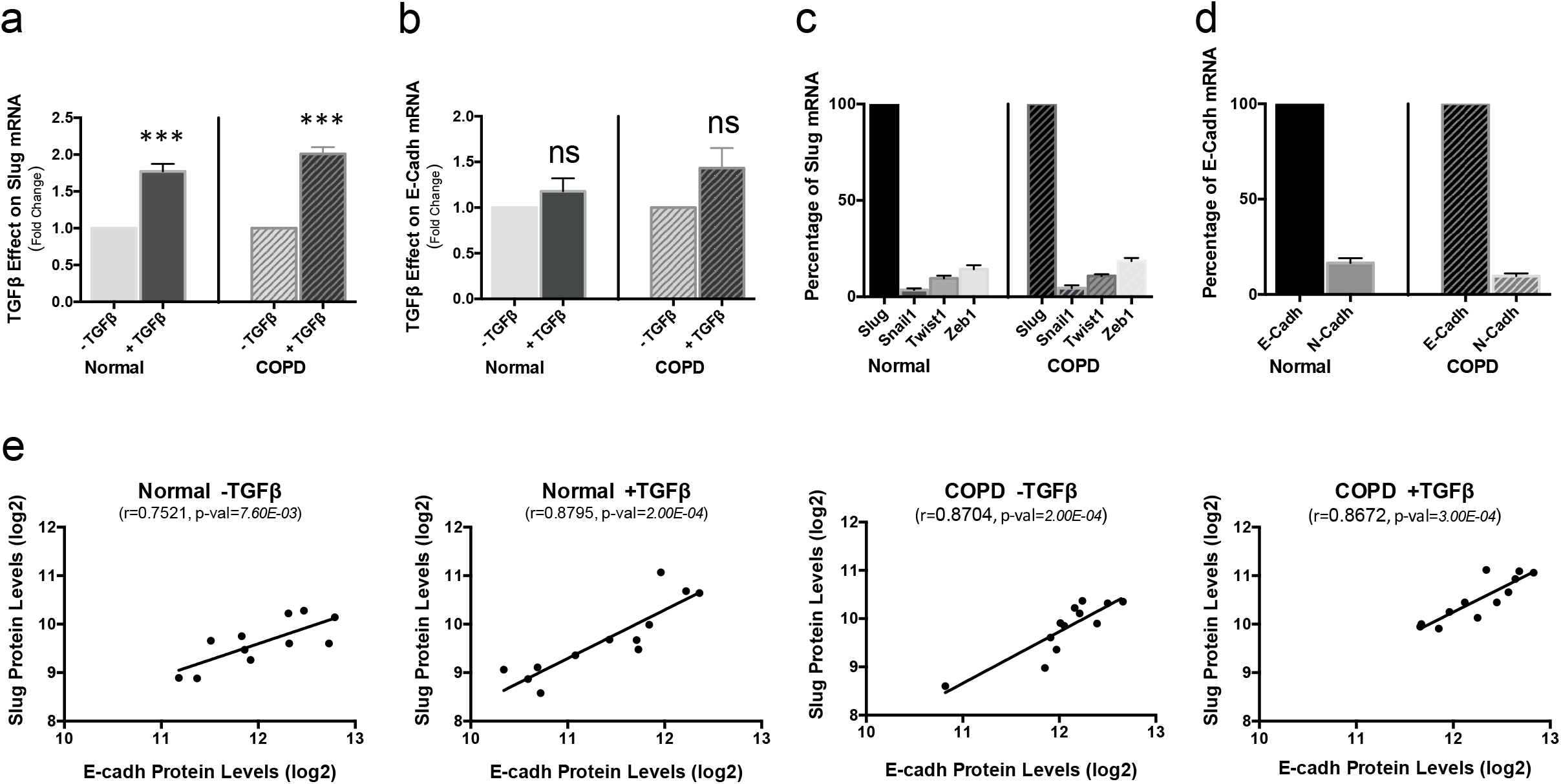
Effect of TGF-β on Slug and EMT-related markers in normal and COPD bronchial epithelial progenitors. Primary bronchial epithelial basal cells, normal and COPD, were grown on filters and at confluence changed to ALI culture to induce differentiation, without TGF-β or in presence of 1ng/ml of TGF-β. Cells were analyzed at day 6 of ALI culture for mRNA (**a-d**) or protein (**e**) expression. (**a-d**) RNA were extracted from normal and COPD cells and analyzed by RT-qPCR. GAPDH was used to normalize cDNA amounts between samples and results were calculated as a ratio on GAPDH. Data shown represent the mean for n ≥ 11, and experiments were done at least in duplicate. (**a, b**) Results are presented as the fold-change induced by TGF-β on mRNA expression of Slug (**a**), E-cad/CDH1 (**b**) with mean ±SEM, and compare normal and COPD cells. (**c**) Expression of the EMT-inducing transcription factors Snail1, Twist1 and Zeb1 shown as a percentage of Slug mRNA, with mean ±SEM. (**d**) Expression of the mesenchymal marker N-cad/CDH2 shown as a percentage of the epithelial marker E-cad/CDH1 mRNA, with mean ±SEM. Statistical significance is at P value < 1.00E-03 *** as indicated. ns: non significant. (**e**) Proteins were extracted from normal and COPD cells and quantified by RPPA. Correlations between Slug and E-cad protein levels in absence or presence of TGF-β for normal and COPD cells were determined. Results are Pearson correlations calculated with log2 (expression levels) and are presented as scatter plots with a regression line. Statistical significance is at P value < 1.00E-02 ** or < 1.00E-03 *** as indicated. ns: non significant.

Moreover when we determined the expression of Slug and E-cad proteins by Reverse Phase Protein Array (RPPA), a technology that allows studying all the samples simultaneously and thus reduces variability due to technical bias [32], we found a strong positive correlation between Slug and E-cad protein levels both in normal and COPD cells, and in absence or presence of TGF-β (Fig. 2e). Immunocytochemistry shows that Slug is expressed in the cell nuclei both in presence and absence of TGF-β. For E-cad on the other hand, TGF-β leads to delocalization from the cell-cell junctions to the cytoplasm both in normal and COPD cells (Fig. S3a-d) In conclusion, in both normal and COPD progenitor cells, induction of differentiation by TGF-β leads to upregulation of Slug and Vim, while E-cad is delocalized from the cell membrane to the cytoplasm.

### Microarray analysis of Slug knockdown identifies genes involved in proliferation that are repressed downstream of Slug in normal but not COPD cells

We found that, in normal and COPD bronchial progenitor cells, Slug coexpresses with epithelial and mesenchymal markers and that it does not repress E-cad, even in presence of TGFβ. To better understand Slug function in these progenitor cells, we wanted to identify the genes downstream of Slug and determine if these genes were similarly altered between normal and COPD cells. It has been reported that the expression level of Slug determines the differentiation status of keratinocytes [3], and we chose a Slug knockdown approach rather than overexpression, in order to avoid unspecific off target binding of the overexpressed protein to irrelevant genes. We knocked down Slug in normal and COPD cells using shRNA that were validated and we selected the most efficient shRNA within the coding strand in order to limit non-specific downstream genes [33]. To determine the genes whose expression was modified by Slug knockdown compared to a control siRNA, we performed a total RNA microarray analysis using an Affymetrix human Gene 2.1 ST GeneChip, that allows to test >30,000 coding transcripts and 11,000 long intergenic non-coding transcripts.

Slug Knockdown resulted in a statistically significant decrease of Slug mRNA with knockdown levels determined by RT-qPCR of 43% (fold-change 1.77, P value <1.00E-04) and 54% (fold-change 2.17, P value 4.20E-03), respectively for normal and COPD cells. Slug protein levels decreased similarly (Fig. S4a, b). We identified 805 genes whose expression was changed by Slug knockdown in normal cells (P value ≤5.00E-02 and absolute fold-change ≥ 1.2). Among these genes, a majority (514/63.9%) were upregulated upon Slug knockdown, i.e. genes repressed downstream of Slug, which is coherent with Slug being a repressor [34-35].

Indeed, we were particularly interested in genes responding to differentiation and TGF-β and the role of Slug herein. To identify these genes, we performed in parallel another microarray analysis on RNA from basal cells at the onset of differentiation, in presence or absence of TGF-β, using undifferentiated cells as control. Table S1 is the list of the 514 genes upregulated in normal cells with their fold change in normal and COPD cells knocked down for Slug as well as their respective response to differentiation or TGF-β. A large majority of these 514 genes (398/77.4%,) respond to differentiation and/or TGF-β with a fold-change ≥1.2, and, among these 193 (48.5%) respond to both differentiation and TGF-β. We classified these 193 genes in 4 groups according to their type of response and Fig. 3a shows histograms representing the number of genes for each group with small bars indicating the group mean fold-increase induced by Slug knockdown in normal (blue) or COPD (red) cells. The large majority of the genes are in the 2 groups of genes repressed by TGF-β. Among these, the group of genes upregulated during differentiation has a similar mean fold-increase in normal and COPD cells. In contrast, the 68 genes in the group downregulated during differentiation have a much higher mean fold-increase in normal than in COPD cells, showing that these genes are more repressed downstream of Slug in normal cells than in COPD cells. A search for enriched gene pathways using KEGG, REACTOME and Gene Ontology (GO) databases revealed that 45 out of these 68 genes are involved in processes related to proliferation or cell cycle. Table 1 is the list of these 45 genes and it shows that, except for 3 (surligned in grey), they are much less or not repressed downstream of Slug in COPD compared to normal cells, with the mean of the fold-change (log2) for all the genes being 0.12 for COPD and 0.56 for normal cells.

**Table 1.**
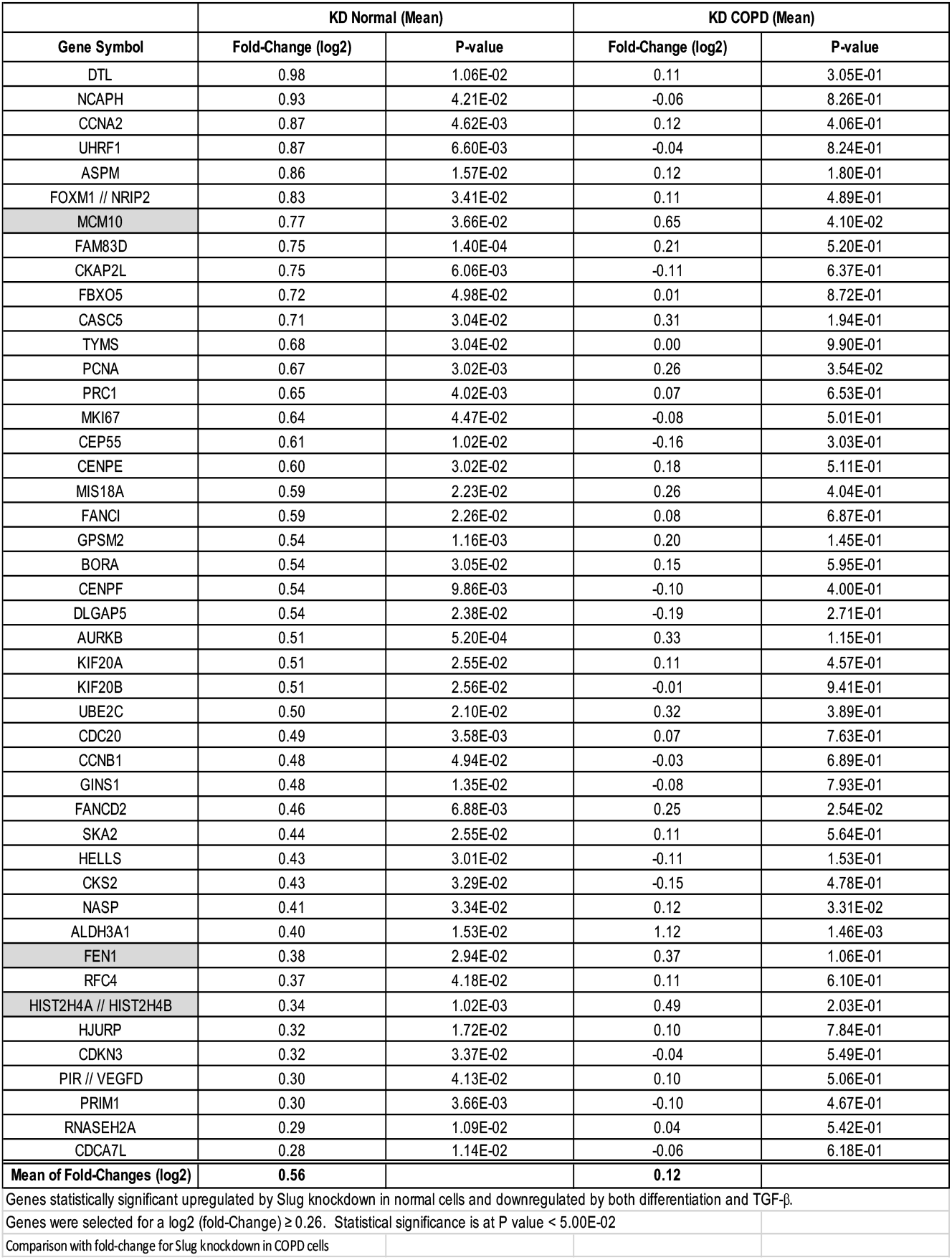
List of 45 proliferation genes upregulated by Slug knockdown in normal bronchial basal/progenitor cells

**Fig. 3.**
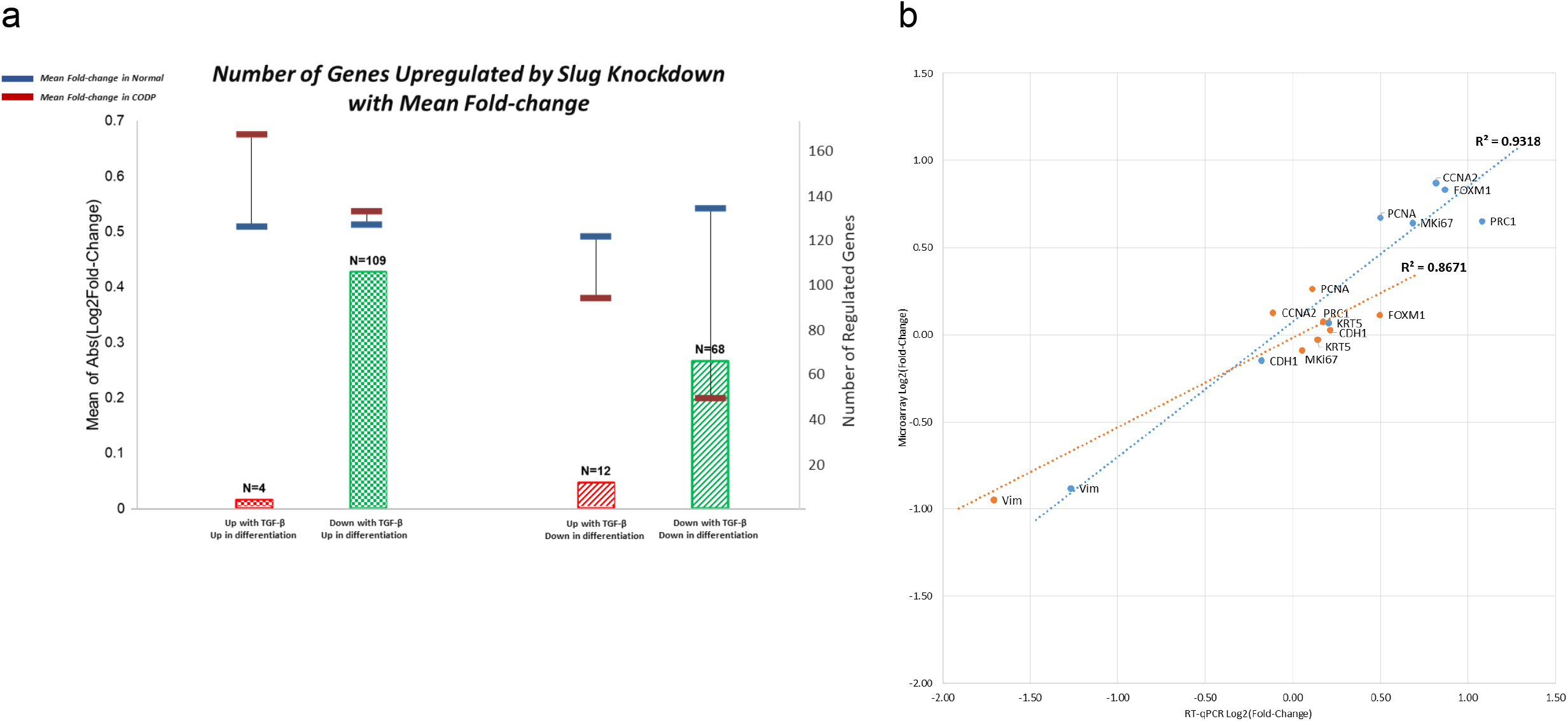
Microarray analysis of Slug knockdown identifies proliferation genes repressed in normal bronchial epithelial progenitors. Knockdown with shRNA specific of Slug were performed on primary bronchial epithelial basal cells, normal and COPD, Knockdown cells were grown on filters and, at confluence, changed to ALI culture to induce differentiation. Cells were analyzed at day 2 of ALI culture for mRNA expression. RNA were extracted from normal and COPD cells and analyzed by microarray analysis on an Affymetrix chip. Data are for n ≥ 3 each, normal and COPD. (**a**) Genes significantly upregulated by Slug knockdown in normal cells and responding to both differentiation and TGF-β were classified in 4 groups according to their response to differentiation and TGF-β. Histograms represent the number of genes in each group with genes upregulated by TGF-β in red and genes downregulated by TGF-β in green, and genes upregulated during differentiation as checkboard and genes downregulated during differentiation as oblique lines. Horizontal bars represent the mean fold-increase for each group of genes, in blue for normal cells and in red for COPD cells. (**b**) Validation of microarray by RT-qPCR and comparison of Slug knockdown effect in normal (blue) and COPD (red) cells for CDH1, KRT5 and VIM genes as well as proliferation-related genes. Results are Pearson correlations calculated with log2 (fold-change) and are presented as scatter plots with regression line and R-square values.

We further studied the expression of 5 of these proliferation-related genes and first confirmed by RT-qPCR the fold change induced by Slug knockdown; we also studied E-cad/CDH1, cytokeratin 5/KRT5 and Vim/VIM genes that were not among the 514 genes significantly increased by Slug knockdown in normal cells. Fig. 3b shows the correlation between microarray and RT-qPCR values for these 8 genes in both normal and COPD cells. The strong correlation (r^2^ values of 0.9318 and 0.8671, respectively for normal and COPD), validates the microarray analysis and also confirms that all 5 proliferation-related genes are upregulated by Slug knockdown in normal but not COPD cells. No significant effect of Slug knockdown is seen on CDH1 and KRT5 expression, while VIM expression decreases significantly and similarly for normal and COPD. This confirms that CDH1/E-cad is not a target of Slug in bronchial progenitor cells from both normal and COPD. KRT5 is also not regulated downstream of Slug, while in contrast, VIM is induced downstream of Slug and this similarly in cells from normal and COPD subjects.

The increased expression of the proliferation-related genes following Slug knockdown observed in normal but not in COPD cells suggests that these genes are directly or indirectly repressed downstream of Slug in normal cells but that this regulation is lost in COPD. This is supported by the strong and statistically significant inverse correlation between the mRNA levels of Slug and these genes observed in undifferentiated normal but not COPD cells. In addition, for the widely used proliferation marker Ki67 [36], a strong inverse correlation also exists between its protein and Slug mRNA levels in normal cells (Table 2 and Fig. 4a).

**Table 2.**
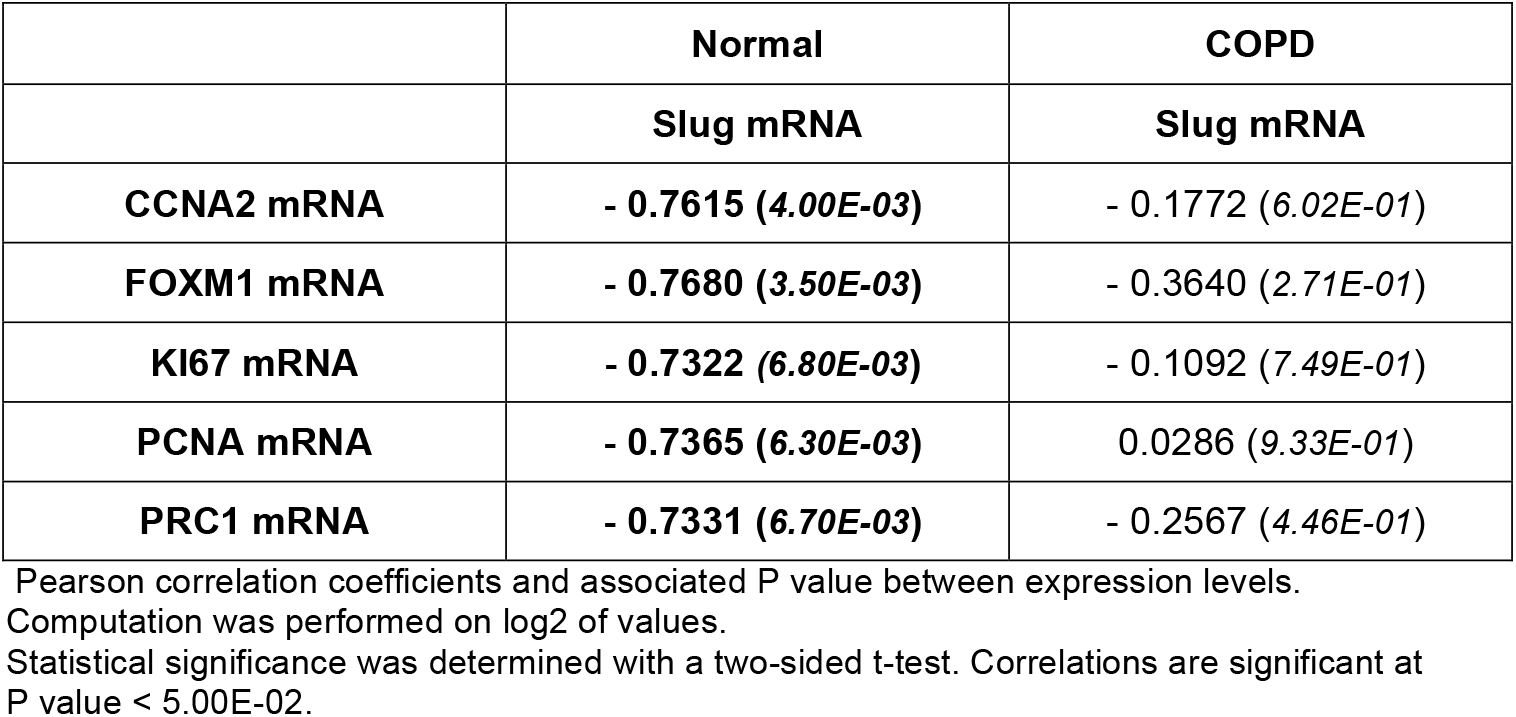
Correlation between proliferation genes and Slug in normal and COPD undifferentiated basal cells

|  | Normal | COPD |
| --- | --- | --- |
|  | Slug mRNA | Slug mRNA |
| **CCNA2 mRNA** | - 0.7615 (4.00E-03) | - 0.1772 (6.02E-01) |
| **FOXM1 mRNA** | - 0.7680 (3.50E-03) | - 0.3640 (2.71E-01) |
| **KI67 mRNA** | - 0.7322 (6.80E-03) | - 0.1092 (7.49E-01) |
| **PCNA mRNA** | - 0.7365 (6.30E-03) | 0.0286 (9.33E-01) |
| **PRC1 mRNA** | - 0.7331 (6.70E-03) | - 0.2567 (4.46E-01) |

Pearson correlation coefficients and associated P value between expression levels.
Computation was performed on log2 of values.
Statistical significance was determined with a two-sided t-test. Correlations are significant at P value < 5.00E-02.

**Fig. 4.**
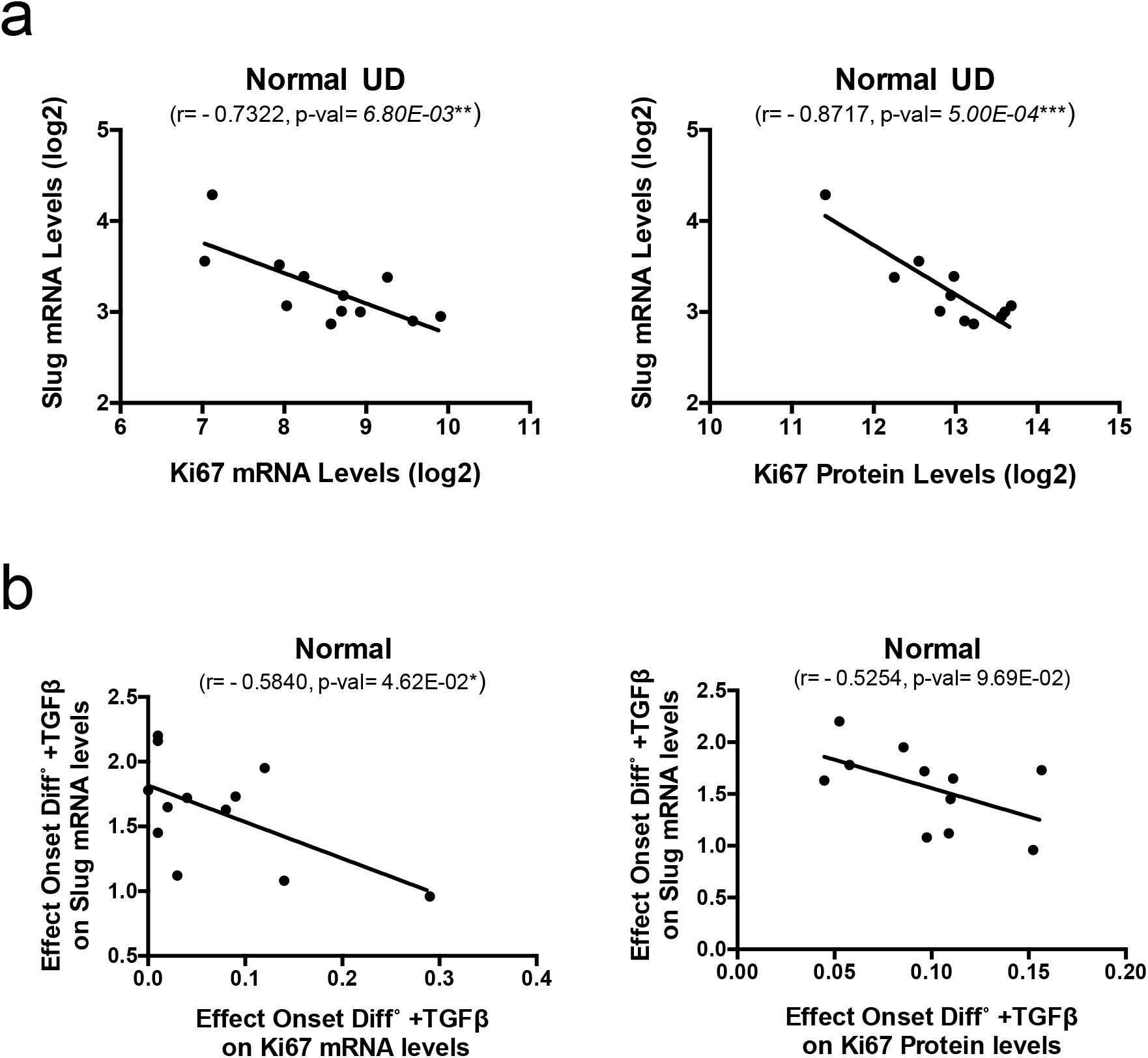
Ki67 and Slug inversely correlate in normal bronchial epithelial progenitors. Normal primary bronchial epithelial basal cells were grown on filters and analyzed undifferentiated at confluence or grown on filters and at confluence changed to ALI culture to induce differentiation, without TGF-β or in presence of 1 ng/ml of TGF-β. Cells were analyzed undifferentiated or at day 6 of ALI culture either for mRNA or for protein expression. RNA or proteins lysates were prepared from normal and COPD cells and analyzed respectively by RT-qPCR and RPPA. For RT-qPCR, GAPDH was used to normalize cDNA amounts between samples and results were calculated as a ratio on GAPDH. Data shown are for n ≥ 11. (**a**) Correlations between Slug mRNA and Ki67 mRNA and protein levels in undifferentiated normal cells. Results are Pearson correlations calculated with log2 (expression levels) and are presented as scatter plots with a regression line. (**b**) Correlations between Ki67, mRNA or protein, and Slug mRNA in normal cells for the effect of differentiation and TGF-β. Results are Pearson correlations calculated with the level of effect and are presented as scatter plots with a regression line. Statistical significance is at P value < 5.00E-02 *, < 1.00E-02 ** or < 1.00E-03 *** as indicated. ns: non significant.

Moreover, we also found, and only in normal cells, significant inverse correlations between the combined effects of differentiation and TGF-β on Slug mRNA levels and that on these proliferation-related gene mRNA levels, except for PCNA. A good tendency for an inverse correlation also exists for Ki67 protein (Table 3 and Fig. 4b).

**Table 3.** Correlation between the effect of differentiation and TGF-β on proliferation genes and on Slug in normal and COPD basal/progenitor cells

|  | Normal | COPD |
| --- | --- | --- |
|  | Slug mRNA | Slug mRNA |
| **CCNA2 mRNA** | - 0.7056 (1.04E-02) | 0.3527 (2.87E-01) |
| **FOXM1 mRNA** | - 0.6003 (3.91E-02) | 0.5383 (8.76E-02) |
| **KI67 mRNA** | - 0.5840 (4.62E-02) | 0.2768 (4.10E-01) |
| **PCNA mRNA** | - 0.2238 (4.84E-01) | 0.2448 (4.68E-01) |
| **PRC1 mRNA** | - 0.6983 (1.15E-02) | 0.4013 (2.21E-01) |

Pearson correlation coefficients and associated P value between the effect of differentiation and TGF-β .
Statistical significance was determined with a two-sided t-test. Correlations are significant at P value < 5.00E-02

### Slug and proliferation-related genes positively correlate in COPD but not normal progenitors during early differentiation when in presence of TGF-β

To futher explore the difference between normal and COPD progenitors for proliferation-related genes that we identified within Slug downstream genes, we determined their expression and compared to the expression of Slug.

In progenitors undifferentiated and at the onset of differentiation without TGF-β, mRNA levels of the proliferation-related genes are not statistically significatively different between normal and COPD progenitors, except for Ki67 mRNA levels that are slightly higher in COPD undifferentiated cells. In contrast at the onset of differentiation if in presence of TGF-β, these genes mRNA levels are higher in COPD. This is statistically significant for CCNA2, FoxM1 and Ki67, and close to significant for PRC1 while PCNA shows only a weak increase that is not statistically significant (Fig. 5a, b). TGF-β decreases the expression of these proliferation-related genes with a mean effect in accordance with the mean difference of their expression levels between normal and COPD cells (Fig. S5).

**Fig. 5.**
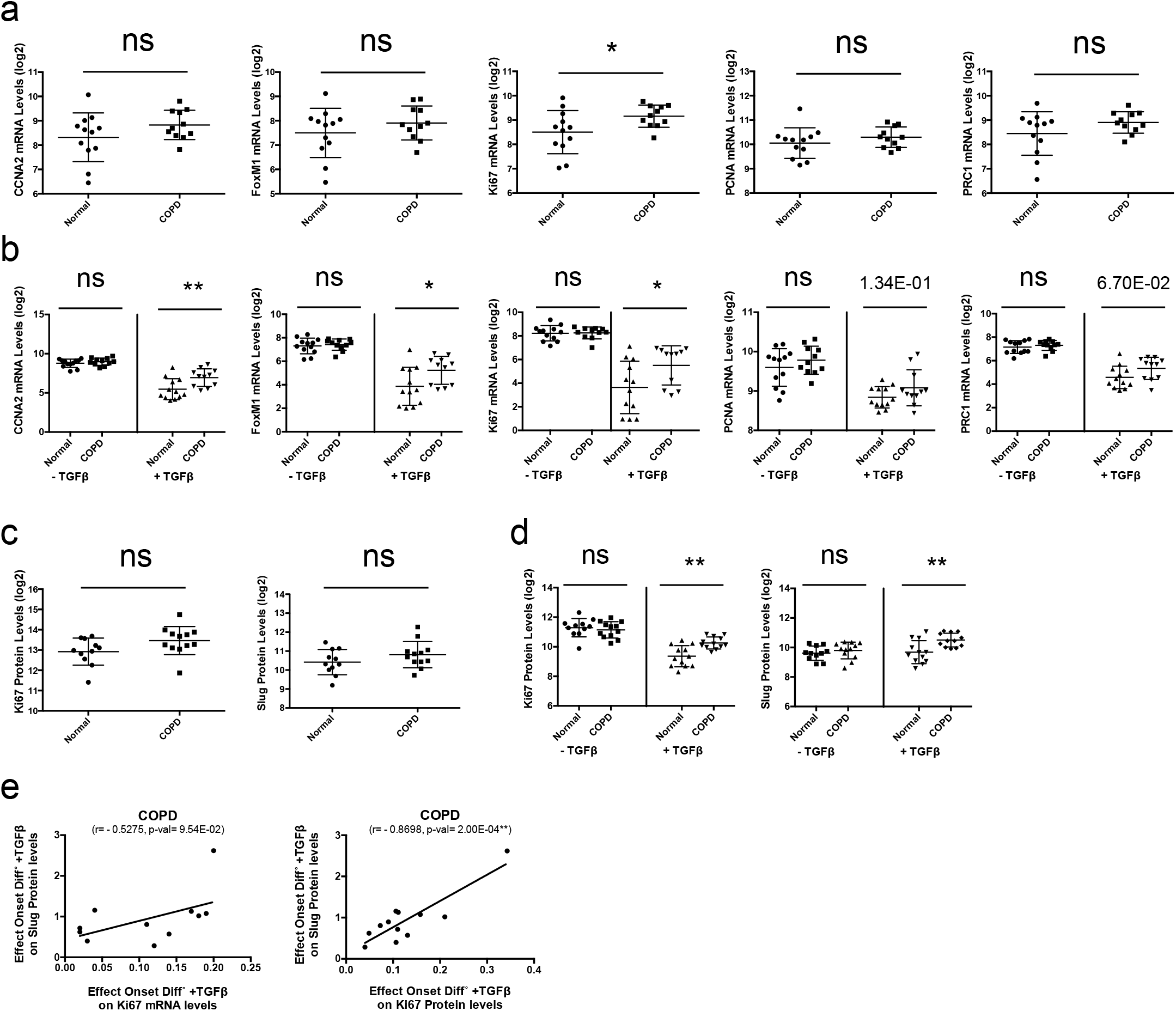
Slug and proliferation genes are more expressed at the onset of differentiation in presence of TGF-β in COPD than in normal bronchial epithelial progenitors. Primary bronchial epithelial basal cells, normal and COPD, were grown on filters and analyzed undifferentiated at confluence or grown on filters and at confluence changed to ALI culture to induce differentiation, without TGF-β or in presence of 1 ng/ml of TGF-β. Cells were analyzed undifferentiated or at day 6 of ALI culture either for mRNA or for protein expression. RNA or proteins lysates were prepared from normal and COPD cells and analyzed respectively by RT-qPCR and RPPA. For RT-qPCR, GAPDH was used to normalize cDNA amounts between samples and results were calculated as a ratio on GAPDH. Data shown are for n ≥ 11. (**a, b**) Comparison of the mRNA levels of proliferation-related genes between normal and COPD cells, undifferentiated (**a**), at the onset of differentiation without TGF-β or in presence of 1 ng/ml of TGF-β (**b**). (**c, d**) Comparison of the protein levels of Ki67 and Slug between normal and COPD cells, undifferentiated (**c**), at the onset of differentiation without TGF-β or in presence of 1 ng/ml of TGF-β (**d**). Results shown are log2 (ratio on GAPDH) for mRNA or log2 (values) for protein and are presented as a scatter plot with the mean ±SD. (**e**) Correlations of the effect of differentiation and TGF-β between Ki67 mRNA or protein, and Slug protein in COPD cells. Results are Pearson correlations calculated with the level of effect and are presented as scatter plots with a regression line. Statistical significance is at P value < 5.00E-02 *, < 1.00E-02 ** or < 1.00E-03 *** as indicated. ns: non significant.

Moreover, we found that Ki67 protein levels in COPD are also not statistically significatively different in undifferentiated progenitors, as well as in progenitors at the onset of differentiation if without TGF-β while if in presence of TGF-β they are significatively higher. Similar results are found for Slug protein (Fig, 5c, d).

We then analyzed the correlation between Slug and proliferation-related genes and found that in COPD but not normal cells, there are positive correlations, significant (FoxM1, PRC1, Ki67 protein) or with a good tendency (CCNA2, Ki67 mRNA), between the combined effects of differentiation and TGF-β on Slug protein levels and that on the proliferation-related gene mRNA and Ki67 protein levels (Table 4 and Fig. 5e). No correlation was found for PCNA gene that was found to have no significatively higher levels in COPD.

**Table 4.**
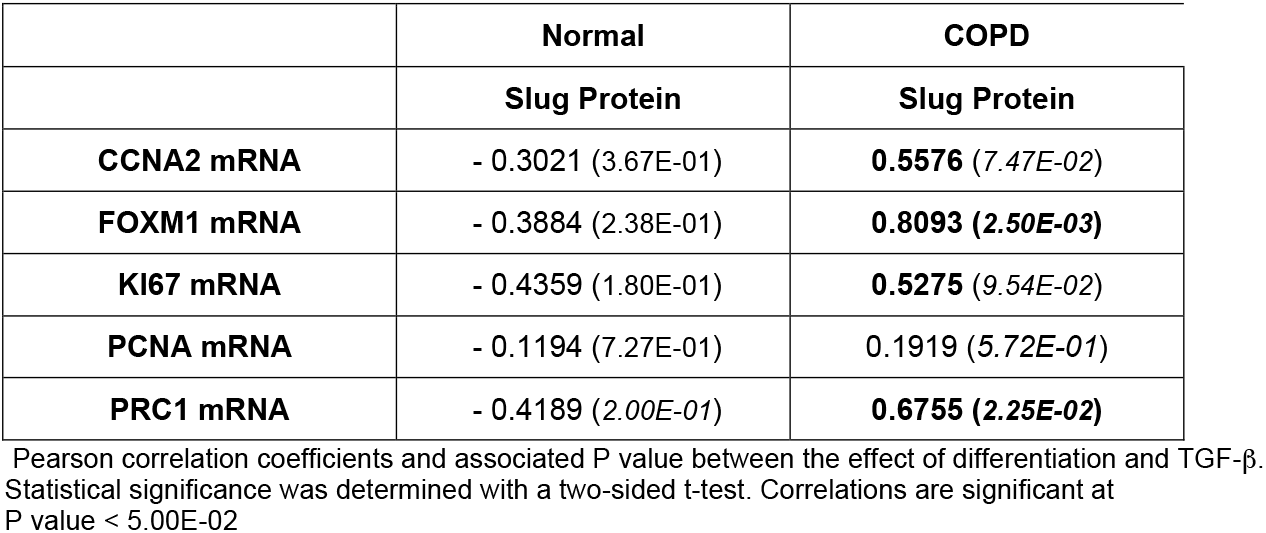
Correlation between the effect of differentiation and TGF-β on proliferation genes and on Slug in normal and COPD basal/progenitor cells

### The higher levels of Slug protein and of proliferation-related genes observed in COPD progenitor cells result from a different combination of TGF-β signaling pathways

We show that Slug and Ki67 protein as well as proliferation-related genes mRNA levels are higher in COPD in early differentiation when in presence of TGF-β. Moreover, we found only in COPD a positive correlation between the combined effect of differentiation and TGF-β on Slug protein level and on proliferation-related gene levels.

However, our Slug knockdown experiment revealed that while in normal cells many proliferation-related genes are repressed downstream of Slug, in COPD these genes are not downstream of Slug.

To find the link between the higher expression of Slug protein and of the proliferation-related genes in COPD progenitor cells in presence of TGF-β, we investigated TGF-β signaling pathways. In addition to the canonical Smad3-dependent TGF-β signaling pathway, we also considered β-catenin (β-cat) signaling pathway known to crosstalk with Smad signaling and to be involved in proliferation [37]. While TGF-β increases the level of phosphorylation of Smad3, i.e. activates Smad3-dependent pathway, both in normal and COPD cells and at similar levels, it turns on β-cat signaling (i.e. decreases phosphorylation of β-cat) only in COPD. In addition, also only in COPD, TGF-β increases the levels of total Smad3 and β-cat (Fig. 6a, b).

**Fig. 6.**
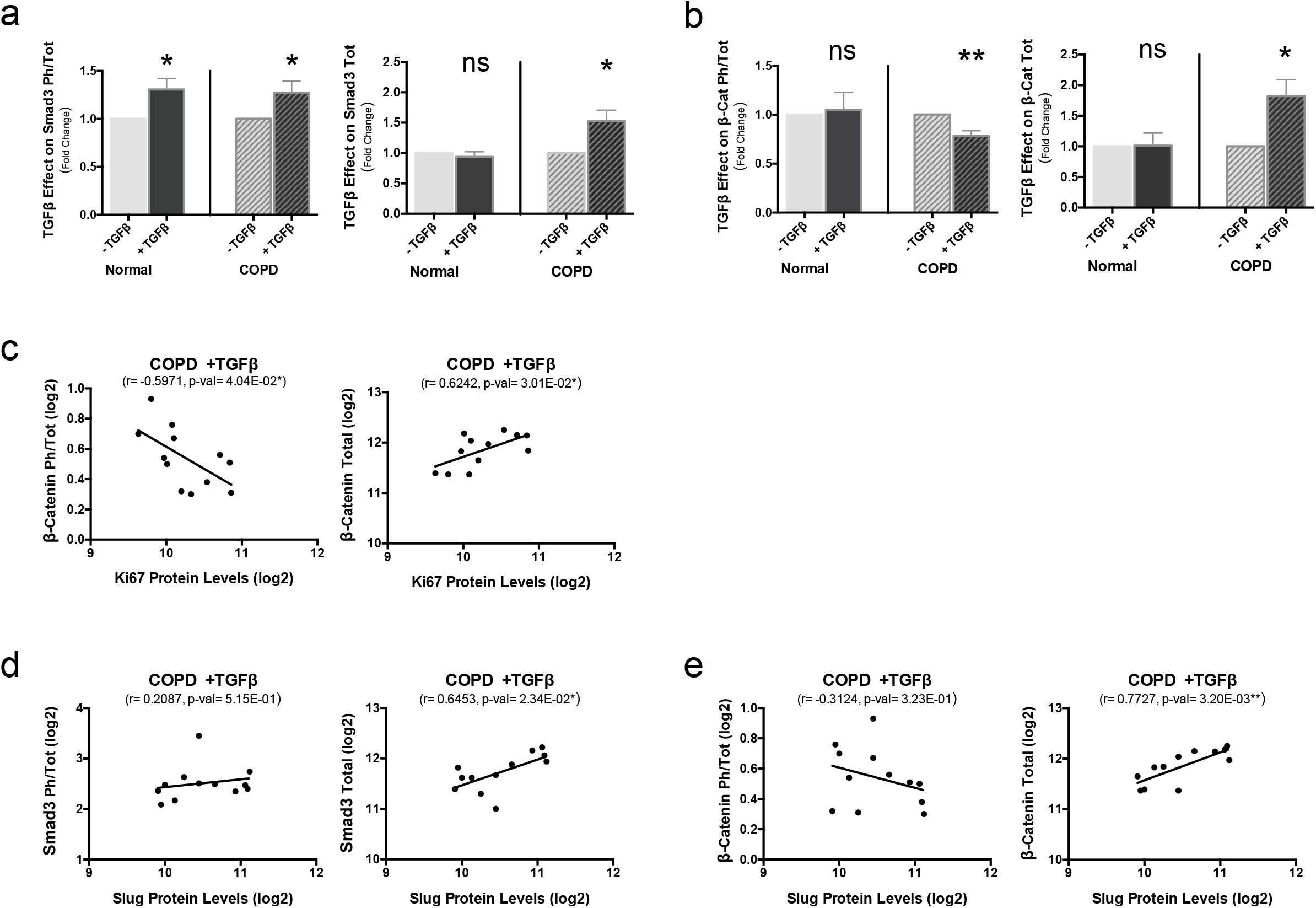
In COPD, TGF-β induced higher levels of Slug protein and proliferation genes through common and different pathways. Primary bronchial epithelial basal cells, normal and COPD, were grown on filters and at confluence changed to ALI culture to induce differentiation, without TGF-β or in presence of 1 ng/ml of TGF-β. Cells were analyzed at day 6 of ALI culture for protein expression. Proteins lysates were prepared from normal and COPD cells and analyzed by RPPA. Data shown are for n ≥ 11. (**a, b**) Comparison of TGF-β effect on the ratio Phospho/Total or total protein expression levels of Smad3 (**a**) and β-cat (b) between normal and COPD cells. Results are presented as the fold-change induced by TGF-β with mean ±SEM. (**c-e**) Correlations between Smad3 or β-cat and Ki67 or Slug protein levels in COPD cells in presence of TGF-β. Ki67 protein levels and β-cat (Total protein level or ratio Phospho/Total) (**c**), Slug protein levels and Smad3 (Total protein level or ratio Phospho/Total) (**d**) or β-cat (Total protein level or ratio Phospho/Total) (**e**). Results are Pearson correlations calculated with log2 (expression levels) or log2 (Ratio Phospho/Total) and are presented as scatter plots with a regression line. Statistical significance is at P value < 5.00E-02 *, < 1.00E-02 ** or < 1.00E-03 *** as indicated. ns: non significant.

In COPD, there is no correlation between proliferation-related genes and Smad3, while good to strong positive correlations are found between the levels of these proliferation-related gene as well as Ki67 protein levels and total β-cat levels. In addition, with the exception of PCNA, positive correlations, significant or as a strong tendency (CCNA2) are found between the activation of β-cat (i.e. decrease of β-cat phosphorylation) and proliferation-related gene levels as well as of Ki67 protein (Table 5, Fig. 6c). On the other hand, Slug protein levels does not correlate with the phosphorylation levels of Smad3 or β-cat while it correlates positively with both total Smad3 and β-cat levels (Fig. 6d, e). This reveals that in COPD cells, the higher levels of Slug protein and of the proliferation-related gene expression result from different TGF-β downstream pathways.

**Table 5.**
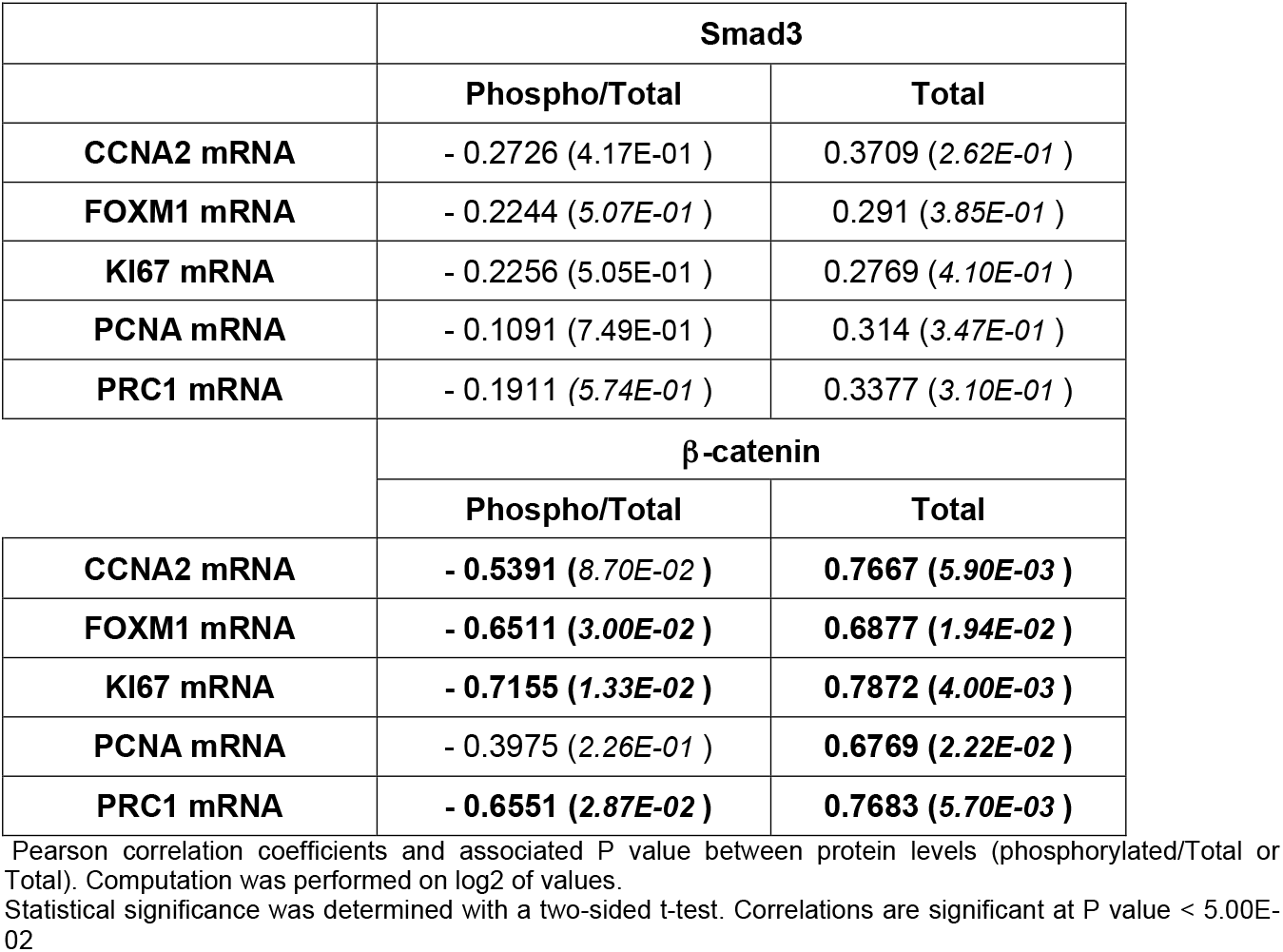
Correlation between the expression levels of proliferation genes and Smad3 or β-cat in COPD basal/progenitor cells in presence of TGF-β.

|  | Smad3 |  |
| --- | --- | --- |
|  | Phospho/Total | Total |
| **CCNA2 mRNA** | - 0.2726 (4.17E-01 ) | 0.3709 (2.62E-01 ) |
| **FOXM1 mRNA** | - 0.2244 (5.07E-01 ) | 0.291 (3.85E-01 ) |
| **KI67 mRNA** | - 0.2256 (5.05E-01 ) | 0.2769 (4.10E-01 ) |
| **PCNA mRNA** | - 0.1091 (7.49E-01 ) | 0.314 (3.47E-01 ) |
| **PRC1 mRNA** | - 0.1911 (5.74E-01 ) | 0.3377 (3.10E-01 ) |
| | β-catenin | |
|  | Phospho/Total | Total |
| **CCNA2 mRNA** | - **0.5391 (8.70E-02 )** | **0.7667 (5.90E-03 )** |
| **FOXM1 mRNA** | - **0.6511 (3.00E-02 )** | **0.6877 (1.94E-02 )** |
| **KI67 mRNA** | - **0.7155 (1.33E-02 )** | **0.7872 (4.00E-03 )** |
| **PCNA mRNA** | - 0.3975 (2.26E-01 ) | **0.6769 (2.22E-02 )** |
| **PRC1 mRNA** | - **0.6551 (2.87E-02 )** | **0.7683 (5.70E-03 )** |

Pearson correlation coefficients and associated P value between protein levels (phosphorylated/Total or Total). Computation was performed on log2 of values. Statistical significance was determined with a two-sided t-test. Correlations are significant at P value < 5.00E-02

## Discussion

In this work, we show that in bronchial basal/progenitor cells, Slug co-expresses with epithelial and mesenchymal markers and, despite being expressed in the cell nuclei, does not repress E-cad, even in presence of TGF-β, and this in normal as well as COPD cells. Using a Slug knockdown approach to identify genes downstream of Slug, we confirm that E-cad is not a target of Slug in normal and COPD progenitors. By selecting among the genes downstream of Slug the one that respond to both differentiation and TGF-β, we reveal that such genes are different between normal and COPD basal/progenitor cells, with in particular genes involved in cell cycle and proliferation that are repressed downstream of Slug in normal but not in COPD. We studied the expression of several of these proliferation-related genes, including the widely used proliferation marker Ki67, and in line with the knockdown results, we show that only in normal progenitors there is strong negative correlations between Slug and the proliferation-related gene mRNA levels and we also find good negative correlations between the effect of TGF-β at the onset of differentiation on Slug and on these proliferation genes. In COPD progenitors, we show that these proliferation-related genes have higher mRNA levels at the onset of differentiation when in presence of TGF-β than in normal cells; in these same conditions Ki67 and Slug protein have also higher levels in COPD, and in addition, there is positive correlations between Slug protein and these proliferation-related genes. While Smad3 is similarly activated by TGF-β in normal and COPD cells, β-cat is activated and both Smad3 and β-cat total protein are increased only in COPD. Slug protein correlate with Smad3 and β-cat total protein levels while proliferation genes correlate with β-cat activated and total protein levels, indicating that Slug and proliferation genes are increased by different TGF-β-induced mechanisms (see graphical abstract).

We report that despite expression of Slug in the nuclei of bronchial basal/progenitors, these cells express both epithelial and mesenchymal markers simultaneously, and this in normal and COPD cells and similarly in presence or absence of TGF-β. In particular, we find that the epithelial cytoskeletal protein KRT5 coexpresses with the mesenchymal cytoskeletal proteins Vim and Acta2 and that the epithelial junctional protein E-cad is well expressed, even in presence of TGF-β. In mammary stem cells co-expression of epithelial and mesenchymal markers, including Slug and E-cad, has also been reported leading to the concept of hybrid phenotype [38-39].

In addition to co-expression of Slug and E-cad proteins, we also find a strong positive correlation between these 2 proteins in normal and COPD progenitors, in absence or presence of TGF-β. Our Slug knockdown reveals that E-cad/CDH1 is not among the genes whose expression is modified by Slug knockdown in normal as well as COPD progenitors, showing that it is not a target of Slug in bronchial basal/progenitor cells. The positive correlations that exist between Slug and E-cad indicates then that these 2 proteins are co-regulated in normal and COPD progenitors and in presence of TGF-β. This is in line with the cooperation of Slug and E-cad that has been reported in mammary stem/progenitor cells [40].

We identified Vim among the genes whose expression is induced downstream of Slug and it is induced at similar levels in normal and COPD cells. These results are in agreement with a report showing that Slug is crucial for the up-regulation of Vim by TGF-β in epithelial cells [19]. COPD cells have been described to reconstitute a bronchial epithelium with many anomalies as well as features of EMT [9]. Several studies have reported features of EMT in COPD bronchial epithelium. However, this is based on studies only revealing cells that double stain for epithelial and mesenchymal markers, including Vim, or on studies showing only a slightly lower expression of E-cad in COPD. Moreover, theses studies concern cells from *ex-vivo* epithelium or fully differentiated epithelium *in vitro* ALI culture, and suggest that COPD cells may be rather imprinted for EMT, and in such, have a higher potential to enter EMT in permissive conditions [9, 41-42]

Our work reveals that the genes downstream of Slug that respond to differentiation and TGF-β are different between normal and COPD cells. In particular we identified a large set of proliferation-related genes that are repressed downstream of Slug in normal but not COPD progenitor cells. We studied several of these genes and show that they are more expressed at the onset of differentiation in presence of TGF-β in COPD when compared to normal progenitors. Moreover, in COPD, in contrast to normal cells, positive correlations are found between the effect of TGF-β on Slug and on the proliferation-related genes. Since in COPD these genes are not dowstream of Slug, these correlations reveal that they are coregulated with Slug by TGF-β. Our results confirm the link between Slug and proliferation previously reported, and can provide an explanation to the fact that some studies report Slug to be a repressor of proliferation [43-45], while others report a positive link with proliferation [2, 46-47]. Normal and COPD basal cells may represent two cell states with a different regulation of proliferation genes and of Slug by TGF-β, and this could reflect what is observed in the different studies that use cells from different origin.

The set of proliferation-related genes studied have slightly different response but among the genes studied, PCNA is apart: It is the gene with the least difference between normal and COPD of Slug KD effect, and this is coherent with the fact that there is no significant difference of expression between normal and COPD and no correlation with Slug for the effect of TGF-β. In addition to its role in DNA replication, PCNA has also a role in DNA repair, making it a poor proliferation marker.Our results on the expression of TGF-β signaling pathways show that progenitor cells from COPD respond differently than normal progenitors to TGF-β; with in particular an increase of total Smad3 and of activated and total β-cat that is not observed in normal cells. The proliferation-related genes expression levels correlate with the levels of activated and total β-cat. β-cat signaling pathway is involved in the regulation of proliferation [37] and we can speculate from our results that in COPD, β-cat signaling is involved in the higher expression of the proliferation genes observed in presence of TGF-β. On the other hand, Slug protein levels correlate with total Smad3 and β-cat and not with the activated form. Slug, Smad3 and β-cat protein have all been reported to be stabilized following GSK-3β inactivation [48-50] and we can speculate that in COPD cells, the increased levels of these proteins results from such a mechanism induced by TGF-β. Overall, our results show that the increase of Slug protein and proliferation-related genes observed in COPD basal/progenitor cells result from different TGF-β.induced mechanisms, and that these mechanisms are not operating in normal cells.

TGF-β function is complex and has a wide spectrum of effects depending on cell state: it is a cell cycle inhibitor in normal cells and a tumor promoter in malignant cells [51-52]. The difference between normal and COPD cells that we report could reflect these antagonistic effects of TGF-β, with COPD cells being in a premalignant state. Moreover, we can speculate that the higher expression of Slug protein seen in COPD progenitor cells in presence of TGF-β could increase during repeated injury. Slug levels of expression define its function [3] and overexpression of Slug induces EMT in epithelial cells [34]. We can also speculate that such deregulations of the progenitor cells could ultimately lead to a shift in Slug function, becoming then an EMT-inducing factor, and this could explain the higher EMT features found in COPD epithelium and the increase risks for COPD to develop lung carcinomas.

## Acknowledgements

We thank the Pulmonary Department, the Pathology Department and the Thoracic Surgery Department at Bichat-Claude Bernard University Hospital (Paris, France) and INSERM UMR 1152 for providing lung tissues and isolating the cells. We thank Audrey Rapinat and David Gentien at the Genomics platform, Curie Institute (Paris, France) for Affymetrix GeneChip hybridization. We thank Dusko Ilic and Pierre Savagner for critical reading of the manuscript and helpful discussions. P.L. is supported by the French National Center for Scientific Research (CNRS). This work was supported by a donation from Association Science et Technologie (Groupe Servier) to P.L. and by funding from French National Institute for Medical Research (INSERM).

## Compliance with ethical standards

### Ethical approval

Human lung tissues were obtained from patients undergoing lung lobectomy for peripheral lung carcinoma after receiving written informed consent. The study was approved by the ethics committee of Paris Nord, *IRB 00006477* Paris 7 University, France.

## Authors’ contibutions

a-Conceived and Designed Experiments: b-Performed the experiments: c-Analyzed the Data: d-Drafted the Article: CBB: a, b, c; CC: b, c; A J: c; BO: b, c ; AC: b, c ; PdlG: c, d; LdK: c, d; PL: a, b, c, d.

## References

1. Nieto, M. A., Huang, R. Y., Jackson, R. A., & Thiery, J. P. (2016). Emt: 2016. Cell, 166(1), 21–45.

2. Nassour, M., Idoux-Gillet, Y., Selmi, A., et al. (2012). Slug controls stem/progenitor cell growth dynamics during mammary gland morphogenesis. PLoS One, 7(12), e53498.

3. Mistry, D. S., Chen, Y., Wang, Y., Zhang, K., & Sen, G. L. (2014). SNAI2 controls the undifferentiated state of human epidermal progenitor cells. Stem Cells, 32(12), 3209–3218.

4. Tesei, A., Zoli, W., Arienti, C., et al. (2009). Isolation of stem/progenitor cells from normal lung tissue of adult humans. Cell Prolif, 42(3), 298–308.

5. Hackett, N. R., Shaykhiev, R., Walters, M. S., et al. (2011). The human airway epithelial basal cell transcriptome. PLoS One, 6(5), e18378.

6. Rock, J. R., Onaitis, M. W., Rawlins, E. L., et al. (2009). Basal cells as stem cells of the mouse trachea and human airway epithelium. Proc Natl Acad Sci U S A, 106(31), 12771–12775.

7. Rock, J. R., Randell, S. H., & Hogan, B. L. (2010). Airway basal stem cells: a perspective on their roles in epithelial homeostasis and remodeling. Dis Model Mech, 3(9-10), 545–556.

8. Fulcher, M. L., Gabriel, S., Burns, K. A., Yankaskas, J. R., & Randell, S. H. (2005). Well-differentiated human airway epithelial cell cultures. Methods Mol Med, 107, 183–206.

9. Gohy, S. T., Hupin, C., Fregimilicka, C., et al. (2015). Imprinting of the COPD airway epithelium for dedifferentiation and mesenchymal transition. Eur Respir J, 45(5), 1258–1272.

10. Mertens, T. C. J., Karmouty-Quintana, H., Taube, C., & Hiemstra, P. S. (2017). Use of airway epithelial cell culture to unravel the pathogenesis and study treatment in obstructive airway diseases. Pulm Pharmacol Ther, 45, 101–113.

11. Rigden, H. M., Alias, A., Havelock, T., et al. (2016). Squamous Metaplasia Is Increased in the Bronchial Epithelium of Smokers with Chronic Obstructive Pulmonary Disease. PLoS One, 11(5), e0156009.

12. Sohal, S. S., & Walters, E. H. (2013). Role of epithelial mesenchymal transition (EMT) in chronic obstructive pulmonary disease (COPD). Respir Res, 14, 120.

13. Steiling, K., van den Berge, M., Hijazi, K., et al. (2013). A dynamic bronchial airway gene expression signature of chronic obstructive pulmonary disease and lung function impairment. Am J Respir Crit Care Med, 187(9), 933–942.

14. Randell, S. H. (2006). Airway epithelial stem cells and the pathophysiology of chronic obstructive pulmonary disease. Proc Am Thorac Soc, 3(8), 718–725.

15. Powell, H. A., Iyen-Omofoman, B., Baldwin, D. R., Hubbard, R. B., & Tata, L. J. (2013). Chronic obstructive pulmonary disease and risk of lung cancer: the importance of smoking and timing of diagnosis. J Thorac Oncol, 8(1), 6–11.

16. Young, R. P., Hopkins, R. J., Christmas, T., Black, P. N., Metcalf, P., & Gamble, G. D. (2009). COPD prevalence is increased in lung cancer, independent of age, sex and smoking history. Eur Respir J, 34(2), 380–386.

17. Mahmood, M. Q., Sohal, S. S., Shukla, S. D., et al. (2015). Epithelial mesenchymal transition in smokers: large versus small airways and relation to airflow obstruction. Int J Chron Obstruct Pulmon Dis, 10, 1515–1524.

18. Mahmood, M. Q., Reid, D., Ward, C., et al. (2017). Transforming growth factor (TGF) beta1 and Smad signalling pathways: A likely key to EMT-associated COPD pathogenesis. Respirology, 22(1), 133–140.

19. Slabakova, E., Pernicova, Z., Slavickova, E., Starsichova, A., Kozubik, A., & Soucek, K. (2011). TGF-beta1-induced EMT of non-transformed prostate hyperplasia cells is characterized by early induction of SNAI2/Slug. Prostate, 71(12), 1332–1343.

20. Xu, J., Lamouille, S., & Derynck, R. (2009). TGF-beta-induced epithelial to mesenchymal transition. Cell Res, 19(2), 156–172.

21. Yumoto, K., Thomas, P. S., Lane, J., et al. (2013). TGF-beta-activated kinase 1 (Tak1) mediates agonist-induced Smad activation and linker region phosphorylation in embryonic craniofacial neural crest-derived cells. J Biol Chem, 288(19), 13467–13480.

22. Bryant, D. M., Datta, A., Rodriguez-Fraticelli, A. E., Peranen, J., Martin-Belmonte, F., & Mostov, K. E. (2010). A molecular network for de novo generation of the apical surface and lumen. Nat Cell Biol, 12(11), 1035–1045.

23. Leroy, P., & Mostov, K. E. (2007). Slug is required for cell survival during partial epithelial-mesenchymal transition of HGF-induced tubulogenesis. Mol Biol Cell, 18(5), 1943–1952.

24. Troncale, S., Barbet, A., Coulibaly, L., et al. (2012). NormaCurve: a SuperCurve-based method that simultaneously quantifies and normalizes reverse phase protein array data. PLoS One, 7(6), e38686.

25. Maubant, S., Tahtouh, T., Brisson, A., et al. (2018). LRP5 regulates the expression of STK40, a new potential target in triple-negative breast cancers. Oncotarget, 9(32), 22586–22604.

26. de la Grange, P., Dutertre, M., Correa, M., & Auboeuf, D. (2007). A new advance in alternative splicing databases: from catalogue to detailed analysis of regulation of expression and function of human alternative splicing variants. BMC Bioinformatics, 8, 180.

27. de la Grange, P., Dutertre, M., Martin, N., & Auboeuf, D. (2005). FAST DB: a website resource for the study of the expression regulation of human gene products. Nucleic Acids Res, 33(13), 4276–4284.

28. de la Grange, P., Gratadou, L., Delord, M., Dutertre, M., & Auboeuf, D. (2010). Splicing factor and exon profiling across human tissues. Nucleic Acids Res, 38(9), 2825–2838.

29. Kanehisa, M., Goto, S., Sato, Y., Furumichi, M., & Tanabe, M. (2012). KEGG for integration and interpretation of large-scale molecular data sets. Nucleic Acids Res, 40(Database issue), D109–114.

30. Haw, R., & Stein, L. (2012). Using the reactome database. Curr Protoc Bioinformatics, Chapter 8, Unit8 7.

31. Huang da, W., Sherman, B. T., & Lempicki, R. A. (2009). Systematic and integrative analysis of large gene lists using DAVID bioinformatics resources. Nat Protoc, 4(1), 44–57.

32. Akbani, R., Becker, K. F., Carragher, N., et al. (2014). Realizing the promise of reverse phase protein arrays for clinical, translational, and basic research: a workshop report: the RPPA (Reverse Phase Protein Array) society. Mol Cell Proteomics, 13(7), 1625–1643.

33. Coussens, M. J., Corman, C., Fischer, A. L., Sago, J., & Swarthout, J. (2011). MISSION LentiPlex pooled shRNA library screening in mammalian cells. J Vis Exp(58).

34. Bolos, V., Peinado, H., Perez-Moreno, M. A., Fraga, M. F., Esteller, M., & Cano, A. (2003). The transcription factor Slug represses E-cadherin expression and induces epithelial to mesenchymal transitions: a comparison with Snail and E47 repressors. J Cell Sci, 116(Pt 3), 499–511.

35. Savagner, P. (2001). Leaving the neighborhood: molecular mechanisms involved during epithelial-mesenchymal transition. Bioessays, 23(10), 912–923.

36. Jurikova, M., Danihel, L., Polak, S., & Varga, I. (2016). Ki67, PCNA, and MCM proteins: Markers of proliferation in the diagnosis of breast cancer. Acta Histochem, 118(5), 544–552.

37. Logan, C. Y., & Nusse, R. (2004). The Wnt signaling pathway in development and disease. Annu Rev Cell Dev Biol, 20, 781–810.

38. Jolly, M. K., Tripathi, S. C., Jia, D., et al. (2016). Stability of the hybrid epithelial/mesenchymal phenotype. Oncotarget, 7(19), 27067–27084.

39. Ye, G. D., Sun, G. B., Jiao, P., et al. (2016). OVOL2, an Inhibitor of WNT Signaling, Reduces Invasive Activities of Human and Mouse Cancer Cells and Is Down-regulated in Human Colorectal Tumors. Gastroenterology, 150(3), 659–671 e616.

40. Sterneck, E., Poria, D. K., & Balamurugan, K. (2020). Slug and E-Cadherin: Stealth Accomplices? Front Mol Biosci, 7, 138.

41. Mahmood, M. Q., Walters, E. H., Shukla, S. D., et al. (2017). beta-catenin, Twist and Snail: Transcriptional regulation of EMT in smokers and COPD, and relation to airflow obstruction. Sci Rep, 7(1), 10832.

42. Sohal, S. S., Reid, D., Soltani, A., et al. (2011). Evaluation of epithelial mesenchymal transition in patients with chronic obstructive pulmonary disease. Respir Res, 12, 130.

43. Sun, Y., Shao, L., Bai, H., Wang, Z. Z., & Wu, W. S. (2010). Slug deficiency enhances self-renewal of hematopoietic stem cells during hematopoietic regeneration. Blood, 115(9), 1709–1717.

44. Turner, F. E., Broad, S., Khanim, F. L., et al. (2006). Slug regulates integrin expression and cell proliferation in human epidermal keratinocytes. J Biol Chem, 281(30), 21321–21331.

45. Wang, W. L., Huang, H. C., Kao, S. H., et al. (2015). Slug is temporally regulated by cyclin E in cell cycle and controls genome stability. Oncogene, 34(9), 1116–1125.

46. Bhat-Nakshatri, P., Appaiah, H., Ballas, C., et al. (2010). SLUG/SNAI2 and tumor necrosis factor generate breast cells with CD44+/CD24-phenotype. BMC Cancer, 10, 411.

47. Phillips, S., Prat, A., Sedic, M., et al. (2014). Cell-state transitions regulated by SLUG are critical for tissue regeneration and tumor initiation. Stem Cell Reports, 2(5), 633–647.

48. Guo, X., Ramirez, A., Waddell, D. S., Li, Z., Liu, X., & Wang, X. F. (2008). Axin and GSK3-control Smad3 protein stability and modulate TGF - signaling. Genes Dev, 22(1), 106–120.

49. Kim, J. Y., Kim, Y. M., Yang, C. H., Cho, S. K., Lee, J. W., & Cho, M. (2012). Functional Regulation of Slug/Snail2 is dependant on GSK-3b-mediated phosphorylation. FEBS J, 279(16), 2929–2939.

50. Liu, C., Li, Y., Semenov, M., et al. (2002). Control of beta-catenin phosphorylation/degradation by a dual mechanism. Cell, 108(6), 837–847.

51. Derynck, R., Akhurst, R. J., & Balmain, A. (2001). TGF-beta signaling in tumor suppression and cancer progression. Nat Genet, 29(2), 117–129.

52. Fynan, T. M., & Reiss, M. (1993). Resistance to inhibition of cell growth by transforming growth factor-beta and its role in oncogenesis. Crit Rev Oncog, 4(5), 493–540.

