## Supplemental Table, figures and appendix for "Proliferation genes repressed by TGF-β are downstream of Slug/Snail2 in normal bronchial epithelial progenitors and are deregulated in COPD"

### S1 Table. List of the 514 genes upregulated by Slug knockdown in normal bronchial basal/progenitor cells

| | KD Normal (Mean) | | KD COPD (Mean) | Diff Normal (Mean) | TGFβ Normal (Mean) |
|---|---|---|---|---|---|
| Gene Symbol | Fold-Change (Log2) | P-value | Fold-Change (Log2) | Fold-Change (Log2) | Fold-Change (Log2) |
| ATP12A | 1.78 | 4.24E-02 | 1.53 | 1.98 | -1.72 |
| B3GNT6 | 1.24 | 1.37E-02 | 0.54 | 0.58 | -0.52 |
| LOC100287834 | 1.21 | 1.58E-02 | -0.34 | -0.48 | 0.50 |
| ENPP4 | 1.21 | 3.95E-02 | 0.86 | 0.34 | -0.45 |
| CLDN10 | 1.16 | 6.50E-03 | 1.18 | 2.99 | 0.04 |
| KIAA1524 | 1.01 | 8.94E-03 | -0.08 | -0.69 | -0.70 |
| POLR1B | 1.00 | 2.82E-02 | 0.24 | -0.10 | -0.48 |
| COX7C | 0.99 | 6.68E-03 | 0.74 | 0.26 | -0.64 |
| CLU // MIR6843 | 0.99 | 4.80E-04 | 2.09 | 0.76 | -1.57 |
| DTL | 0.98 | 1.06E-02 | 0.11 | -0.91 | -1.39 |
| ASF1B | 0.94 | 1.80E-02 | 0.23 | -0.99 | Not Detected |
| NCAPH | 0.93 | 4.21E-02 | -0.06 | -0.44 | -1.11 |
| CAPN5 | 0.93 | 3.99E-02 | 1.38 | 1.36 | -0.89 |
| CCNA2 | 0.87 | 4.62E-03 | 0.12 | -1.16 | -1.24 |
| UHRF1 | 0.87 | 6.60E-03 | -0.04 | -0.85 | -0.70 |
| ASPM | 0.86 | 1.57E-02 | 0.12 | -0.46 | -1.16 |
| NCMAP | 0.84 | 1.76E-03 | Not Detected | 0.68 | 0.04 |
| FAM111B | 0.83 | 2.80E-03 | 0.41 | -0.79 | -1.18 |
| FOXM1 // NRIP2 | 0.83 | 3.41E-02 | 0.11 | -1.42 | -0.90 |
| RAET1E | 0.79 | 2.08E-02 | 0.33 | 0.01 | -0.58 |
| FAM185A | 0.78 | 2.33E-02 | 0.46 | 0.51 | -0.28 |
| BTBD10 | 0.78 | 9.04E-03 | -0.43 | 0.16 | -0.06 |
| DEPDC1 | 0.77 | 3.14E-02 | -0.59 | -1.15 | -1.13 |
| MCM10 | 0.77 | 3.66E-02 | 0.65 | -1.58 | -1.01 |
| PSCA | 0.77 | 1.97E-02 | 1.41 | 1.94 | -1.12 |
| PTPRZ1 | 0.77 | 4.14E-02 | Not Detected | 1.51 | -1.46 |
| CKAP2L | 0.75 | 6.06E-03 | -0.11 | -0.88 | -1.28 |
| FAM83D | 0.75 | 1.40E-04 | 0.21 | -0.52 | -0.98 |
| HMMR | 0.74 | 3.38E-02 | -0.07 | -0.81 | -1.46 |
| ALDH1A1 | 0.74 | 3.31E-02 | 1.60 | 2.27 | -2.40 |
| TMEM45B | 0.74 | 1.60E-04 | 1.32 | 1.17 | -1.83 |
| DGAT2 | 0.74 | 2.22E-02 | 0.76 | 0.96 | -0.57 |
| MBOAT1 | 0.73 | 4.39E-02 | 0.90 | 1.16 | -0.99 |
| FBXO5 | 0.72 | 4.98E-02 | 0.01 | -0.33 | -0.58 |
| KRT80 | 0.72 | 3.05E-02 | 1.21 | -0.54 | 0.42 |
| CYP4B1 | 0.71 | 3.01E-02 | Not Detected | 2.68 | -2.90 |
| CASC5 | 0.71 | 3.04E-02 | 0.31 | -0.99 | -0.80 |
| TIMM8B | 0.71 | 5.34E-03 | -0.37 | 0.19 | -0.66 |
| VTCN1 | 0.71 | 1.73E-02 | 1.09 | 2.50 | -0.65 |
| ATP10B | 0.71 | 3.09E-02 | 0.53 | 1.61 | -0.70 |
| EPHX3 | 0.71 | 1.75E-02 | 0.23 | 0.01 | 0.26 |
| CYP26A1 | 0.70 | 4.86E-02 | 1.48 | 1.62 | -1.58 |
| SGK1 | 0.70 | 2.58E-02 | 0.69 | 1.79 | -0.19 |
| DNAJB6 | 0.70 | 2.88E-02 | 0.04 | -0.23 | 0.86 |
| MIR1299 | 0.70 | 1.01E-02 | 0.34 | 0.06 | 0.45 |
| RPS6KA4 | 0.70 | 3.44E-02 | 0.06 | -0.72 | Not Detected |
| EIF2B3 | 0.69 | 2.83E-02 | -0.16 | -0.11 | -0.52 |
| NPIPB8 | 0.69 | 9.30E-03 | -0.12 | Not Detected | 0.36 |
| PADI3 | 0.68 | 3.35E-02 | 0.52 | -1.82 | -1.14 |
| ST6GALNAC1 | 0.68 | 4.31E-02 | 1.15 | 0.97 | -0.71 |
| MYLIP | 0.68 | 3.06E-02 | 1.00 | 0.82 | -0.82 |
| TYMS | 0.68 | 3.04E-02 | 0.00 | -1.54 | -0.30 |
| CXXC5 | 0.68 | 4.84E-02 | 0.58 | 0.71 | -0.36 |
| MMP13 | 0.68 | 1.21E-02 | 0.63 | 2.73 | -0.01 |
| G085345_T361310 | 0.68 | 3.82E-02 | 0.26 | Not Detected | Not Detected |
| PCNA | 0.67 | 3.02E-03 | 0.26 | -0.53 | -0.43 |
| SLC25A25 | 0.67 | 8.52E-03 | 0.60 | 0.12 | -0.31 |
| ATXN7L1 | 0.67 | 4.75E-02 | -0.82 | -0.08 | 0.25 |
| ZNF644 | 0.67 | 1.36E-02 | 0.31 | 0.40 | 0.04 |
| IKZF2 | 0.66 | 1.61E-02 | 0.51 | 0.44 | -0.52 |
| GPR160 | 0.66 | 5.62E-03 | 1.08 | 0.89 | -0.18 |
| TSPAN6 | 0.66 | 3.11E-02 | 0.95 | 0.51 | -0.16 |
| PRC1 | 0.65 | 4.02E-03 | 0.07 | -0.59 | -1.52 |
| TCN2 | 0.65 | 1.30E-02 | 0.91 | 1.73 | -0.72 |
| RPSAP58 | 0.65 | 2.34E-02 | 0.43 | 0.97 | 0.01 |
| MKI67 | 0.64 | 4.47E-02 | -0.08 | -0.86 | -1.46 |
| CPNE4 | 0.64 | 1.64E-02 | 0.42 | 1.53 | -0.12 |
| B4GALT5 | 0.64 | 1.18E-02 | 0.74 | 0.69 | 0.11 |
| ARRB1 | 0.64 | 2.46E-02 | 0.87 | 0.77 | 0.06 |
| FAM46C | 0.63 | 2.96E-02 | 0.82 | 0.86 | -0.81 |
| ZWILCH | 0.63 | 4.61E-02 | 0.08 | -0.64 | -0.11 |
| SEC11C | 0.63 | 4.81E-02 | 0.33 | -0.45 | -0.04 |
| SYTL2 | 0.63 | 3.28E-03 | 0.69 | 0.63 | 0.19 |
| MAN1C1 | 0.62 | 3.08E-02 | 1.21 | 2.47 | -1.42 |
| LMNB1 | 0.62 | 4.84E-02 | 0.03 | -0.71 | -0.97 |
| BCAS1 | 0.62 | 2.94E-03 | 0.93 | 0.66 | -0.39 |
| CRYL1 | 0.62 | 2.61E-02 | 0.38 | 0.32 | -0.20 |
| CEP55 | 0.61 | 1.02E-02 | -0.16 | -0.96 | -1.52 |
| CERS3 | 0.61 | 3.91E-02 | -0.08 | 0.25 | -0.46 |
| CIR1 | 0.61 | 2.14E-02 | 0.25 | 0.46 | -0.42 |
| PYM1 | 0.61 | 1.13E-02 | 0.24 | -0.29 | -0.12 |
| NME5 | 0.61 | 2.62E-02 | 0.71 | 0.77 | -0.18 |
| AREG | 0.61 | 7.42E-03 | 0.45 | -1.67 | 0.76 |
| KIFC1 | 0.61 | 1.89E-02 | 0.18 | -0.53 | Not Detected |
| GPATCH2 | 0.61 | 3.79E-02 | 0.04 | 0.16 | 0.06 |
| CENPE | 0.60 | 3.02E-02 | 0.18 | -0.72 | -1.24 |
| DEPDC1B | 0.60 | 2.97E-02 | -0.14 | -0.81 | -0.96 |
| SLC25A33 | 0.60 | 8.92E-03 | 0.40 | 0.26 | -0.34 |
| RDH10 | 0.60 | 6.00E-05 | 1.01 | 0.82 | -0.29 |
| NUP54 | 0.60 | 4.96E-02 | -0.01 | -0.19 | -0.19 |
| RBBP5 | 0.60 | 6.98E-03 | 0.23 | 0.28 | -0.19 |
| FEM1A | 0.60 | 4.43E-02 | 0.59 | -0.12 | 0.01 |
| FANCI | 0.59 | 2.26E-02 | 0.08 | -0.66 | -0.74 |
| UPK1B | 0.59 | 4.03E-02 | 0.40 | 0.21 | -0.66 |
| DDAH1 | 0.59 | 4.59E-02 | 0.20 | 0.14 | -0.60 |
| NUCB2 | 0.59 | 1.74E-02 | 0.38 | 0.52 | -0.62 |
| MIS18A | 0.59 | 2.23E-02 | 0.26 | -0.53 | -0.31 |
| MIF4GD | 0.59 | 2.06E-02 | 0.38 | -0.04 | 0.03 |
| ZNF189 | 0.59 | 3.67E-02 | 0.30 | Not Detected | 0.06 |
| LRRC8B | 0.58 | 2.96E-02 | 0.60 | 0.45 | -0.56 |
| DLEU2 | 0.58 | 4.67E-02 | 0.26 | 0.23 | -0.37 |
| PLLP | 0.58 | 4.54E-02 | 0.82 | 0.19 | -0.72 |
| PPM1B | 0.58 | 4.27E-02 | 0.29 | 0.42 | -0.44 |
| RASGEF1B | 0.58 | 1.56E-03 | 0.93 | 0.71 | 0.41 |
| DANCR | 0.57 | 3.63E-02 | 0.26 | -0.60 | -0.49 |
| ANP32B | 0.57 | 4.36E-02 | -0.14 | -0.21 | 0.14 |
| KIF13B | 0.57 | 1.66E-02 | 0.72 | 1.14 | 0.06 |
| UBE2D4 | 0.57 | 8.34E-03 | 0.59 | Not Detected | Not Detected |
| EFCAB11 | 0.56 | 8.50E-03 | 0.31 | 0.45 | -0.58 |
| MTIF2 | 0.56 | 2.96E-03 | 0.14 | 0.29 | -0.51 |
| FOXQ1 | 0.55 | 4.60E-02 | 0.31 | 0.33 | -1.08 |
| SAE1 | 0.55 | 1.52E-02 | 0.21 | 0.15 | -0.36 |
| SLC25A40 | 0.55 | 1.16E-02 | 0.40 | 0.83 | -0.57 |
| PPP6R2 | 0.55 | 2.44E-02 | 0.11 | 0.21 | -0.01 |
| ZDHHC12 | 0.55 | 8.34E-03 | 0.18 | 0.08 | -0.11 |
| DLGAP5 | 0.54 | 2.38E-02 | -0.19 | -0.59 | -1.82 |
| CENPF | 0.54 | 9.86E-03 | -0.10 | -0.59 | -1.49 |
| GPSM2 | 0.54 | 1.16E-03 | 0.20 | -0.45 | -0.63 |
| BORA | 0.54 | 3.05E-02 | 0.15 | -0.37 | -0.73 |
| ITGB3BP | 0.54 | 2.58E-02 | 0.00 | -0.14 | -0.29 |
| MED8 | 0.54 | 4.82E-03 | 0.24 | 0.00 | -0.30 |
| GYG1 | 0.54 | 2.56E-03 | 0.15 | 0.04 | -0.43 |
| GCNT3 | 0.54 | 3.82E-02 | 0.38 | 0.74 | -0.51 |
| SLC12A6 | 0.54 | 4.34E-03 | -0.03 | 0.46 | -0.03 |
| ZMAT2 | 0.54 | 2.79E-02 | 0.29 | 0.19 | -0.15 |
| SEC13 | 0.54 | 1.82E-03 | 0.01 | -0.54 | 0.08 |
| PPP1R8 | 0.54 | 1.73E-02 | 0.16 | -0.07 | 0.00 |
| ZMPSTE24 | 0.54 | 2.68E-02 | 0.10 | 0.00 | 0.16 |
| HID1 | 0.54 | 2.95E-02 | 0.72 | 0.58 | 0.39 |
| FAM134B | 0.54 | 4.53E-02 | 0.76 | Not Detected | Not Detected |
| HSD17B3 | 0.54 | 2.14E-03 | Not Detected | Not Detected | Not Detected |
| CLDN8 | 0.53 | 3.18E-02 | 0.85 | 1.63 | -1.82 |
| UCHL5 | 0.53 | 2.34E-02 | -0.06 | -0.23 | -0.65 |
| PIGR | 0.53 | 4.19E-02 | 0.91 | 3.76 | -0.62 |
| SH3BGRL2 | 0.53 | 1.92E-02 | 0.75 | 1.06 | -0.76 |
| ANXA10 | 0.53 | 4.20E-02 | 0.10 | 0.14 | -0.96 |
| AQR | 0.53 | 1.49E-02 | 0.31 | 0.11 | -0.14 |
| DA063652 | 0.53 | 3.80E-04 | 0.39 | -0.34 | 0.16 |
| CARD11 | 0.53 | 2.45E-02 | 0.23 | -0.21 | 0.76 |
| MEMO1 | 0.52 | 4.00E-05 | -0.16 | -0.43 | -0.34 |
| LRRC8D | 0.52 | 2.73E-02 | 0.20 | 0.33 | -0.44 |
| ALKBH3 // SEC14L1P1 | 0.52 | 2.95E-02 | 0.18 | -0.33 | -0.03 |
| CWF19L2 | 0.52 | 2.03E-02 | 0.30 | -0.14 | -0.12 |
| DOPEY2 | 0.52 | 2.14E-02 | 0.45 | 0.30 | 0.29 |
| YIPF3 | 0.52 | 2.12E-02 | 0.19 | 0.41 | 0.01 |
| RIPK1 | 0.52 | 2.15E-02 | 0.51 | 0.26 | 0.11 |
| KIF20A | 0.51 | 2.55E-02 | 0.11 | -0.88 | -1.58 |
| KIF20B | 0.51 | 2.56E-02 | -0.01 | -0.46 | -1.23 |
| TMTC1 | 0.51 | 1.36E-02 | 0.43 | 1.04 | -1.17 |
| AURKB | 0.51 | 5.20E-04 | 0.33 | -0.96 | -0.99 |
| ELOVL6 | 0.51 | 3.55E-02 | -0.01 | -0.83 | -0.44 |
| TGS1 | 0.51 | 9.88E-03 | 0.40 | 0.10 | -0.29 |
| FLG-AS1 | 0.51 | 9.14E-03 | 0.29 | Not Detected | Not Detected |
| IL20RA | 0.50 | 3.64E-02 | 0.14 | 0.90 | -1.01 |
| TSPAN1 | 0.50 | 3.37E-02 | -0.14 | 0.48 | -0.64 |
| UBE2C | 0.50 | 2.10E-02 | 0.32 | -0.88 | -0.54 |
| PIGK | 0.50 | 2.91E-02 | -0.01 | 0.38 | -0.42 |
| SRPRB | 0.50 | 1.04E-02 | 0.10 | 0.23 | -0.37 |
| PLEKHA7 | 0.50 | 2.14E-02 | 0.65 | 0.32 | -0.49 |
| CYB5A | 0.50 | 1.52E-02 | 0.55 | 0.58 | -0.33 |
| SLIRP | 0.50 | 3.88E-03 | -0.10 | -0.48 | -0.06 |
| CAAP1 | 0.50 | 1.19E-02 | 0.21 | -0.28 | -0.20 |
| PEX11B | 0.50 | 3.78E-02 | 0.45 | 0.11 | -0.16 |
| SAFB2 | 0.50 | 1.33E-02 | 0.19 | 0.24 | 0.48 |
| LINC01572 | 0.50 | 8.16E-03 | Not Detected | Not Detected | 0.21 |
| CDC20 | 0.49 | 3.58E-03 | 0.07 | -1.23 | -1.63 |
| FAM111A | 0.49 | 4.62E-02 | -0.41 | -0.08 | -0.58 |
| SPDEF | 0.49 | 4.54E-03 | 1.00 | 0.52 | -0.59 |
| ERI1 | 0.49 | 3.80E-04 | 0.04 | -0.45 | -0.40 |
| CAB39L | 0.49 | 2.93E-02 | 0.43 | -0.12 | -0.52 |
| CMAS | 0.49 | 2.71E-02 | 0.06 | 0.29 | -0.32 |
| HEATR3 | 0.49 | 3.01E-02 | 0.16 | 0.30 | -0.42 |

| | KD Normal (Mean) | | KD COPD (Mean) | Diff Normal (Mean) | TGFβ Normal (Mean) |
|---|---|---|---|---|---|
| Gene Symbol | Fold-Change (Log2) | P-value | Fold-Change (Log2) | Fold-Change (Log2) | Fold-Change (Log2) |
| PDXDC1 | 0.49 | 4.86E-02 | 0.20 | 0.62 | -0.56 |
| DCLRE1A | 0.49 | 4.40E-02 | 0.37 | 0.06 | -0.28 |
| NRARP | 0.49 | 4.26E-02 | 0.69 | 0.87 | -0.30 |
| GTF2E2 | 0.49 | 4.07E-02 | 0.39 | -0.34 | -0.04 |
| HSPE1 // HSPE1-MOB4 // M | 0.49 | 1.81E-02 | 0.18 | 0.00 | -0.04 |
| DYNC2LI1 | 0.49 | 1.14E-03 | 0.29 | 0.04 | -0.16 |
| GRB2 | 0.49 | 2.85E-02 | 0.32 | 0.12 | 0.10 |
| PLAC8 | 0.48 | 4.74E-03 | 0.46 | 0.78 | -1.79 |
| GINS1 | 0.48 | 1.35E-02 | -0.08 | -1.12 | -0.93 |
| CCNB1 | 0.48 | 4.94E-02 | -0.03 | -1.16 | -0.81 |
| NSF // NSFP1 | 0.48 | 1.25E-02 | 0.30 | 0.66 | -0.50 |
| CEP83 | 0.48 | 4.46E-02 | 0.10 | -0.19 | -0.12 |
| RSPRY1 | 0.48 | 5.52E-03 | 0.06 | 0.18 | -0.12 |
| PTPRM | 0.48 | 3.19E-02 | 0.56 | 0.65 | -0.03 |
| USP38 | 0.48 | 7.72E-03 | 0.06 | 0.18 | 0.07 |
| SLC12A7 | 0.48 | 4.05E-02 | 0.64 | 0.90 | 0.04 |
| NPNT | 0.46 | 2.92E-02 | 0.58 | 1.62 | -1.12 |
| LINC01344 | 0.46 | 9.58E-03 | 0.29 | 1.93 | -1.09 |
| FANCD2 | 0.46 | 6.88E-03 | 0.25 | -0.33 | -0.70 |
| PBX1 | 0.46 | 1.79E-02 | 0.61 | 1.18 | -0.77 |
| AKR1A1 | 0.46 | 2.40E-02 | 0.50 | 0.38 | -0.74 |
| SLC16A9 | 0.46 | 3.56E-02 | 0.24 | 1.08 | -0.99 |
| C14orf2 | 0.46 | 4.07E-02 | 0.08 | -0.04 | -0.42 |
| RPN2 | 0.46 | 4.58E-03 | -0.04 | 0.19 | -0.11 |
| G046774_T202565 | 0.46 | 2.15E-02 | -0.14 | Not Detected | -0.10 |
| CHMP2B | 0.46 | 1.64E-02 | 0.10 | 0.04 | -0.07 |
| ABHD17B | 0.46 | 4.67E-02 | 0.32 | 0.28 | -0.20 |
| MTX3 | 0.46 | 3.21E-02 | -0.20 | -0.21 | 0.03 |
| G088103_T373456 | 0.46 | 4.02E-03 | 0.11 | Not Detected | Not Detected |
| HIBCH | 0.45 | 3.84E-02 | 0.10 | 0.75 | -1.14 |
| SLC34A2 | 0.45 | 1.41E-02 | 0.70 | 4.28 | -1.09 |
| RABGGTB | 0.45 | 4.59E-02 | -0.28 | -0.01 | -0.40 |
| GNL3 | 0.45 | 6.02E-03 | 0.00 | -0.10 | -0.26 |
| MOCS2 | 0.45 | 3.17E-02 | 0.24 | 0.06 | -0.42 |
| HNRNPAB | 0.45 | 1.20E-02 | 0.10 | 0.03 | -0.20 |
| G061649_T266562 | 0.45 | 1.28E-02 | 0.29 | -1.19 | 0.69 |
| DDHD1 | 0.45 | 3.60E-02 | 0.50 | -0.06 | 0.01 |
| AL832650 | 0.45 | 3.91E-02 | -0.30 | 0.63 | 0.00 |
| SNX16 | 0.45 | 1.53E-02 | 0.56 | 0.63 | 0.20 |
| PLGRKT | 0.45 | 1.83E-02 | 0.10 | 0.33 | 0.06 |
| DMRTA1 | 0.45 | 1.99E-02 | 0.29 | 0.08 | 0.04 |
| SKA2 | 0.44 | 2.55E-02 | 0.11 | -0.28 | -0.93 |
| SEMA3A | 0.44 | 2.40E-02 | -0.03 | -0.01 | -0.93 |
| MMS22L | 0.44 | 3.07E-02 | 0.26 | -0.51 | -0.46 |
| WDFY2 | 0.44 | 9.78E-03 | 0.40 | 0.01 | -0.36 |
| KLHL15 | 0.44 | 2.45E-02 | 0.06 | 0.52 | -0.51 |
| DCAF11 | 0.44 | 4.62E-02 | 0.42 | 0.75 | -0.44 |
| MCU | 0.44 | 8.84E-03 | 0.19 | 0.19 | -0.49 |
| ODC1 | 0.44 | 3.50E-02 | 0.08 | -1.53 | 0.00 |
| MORC4 | 0.44 | 2.03E-02 | 0.21 | 0.00 | -0.10 |
| ANKRD40 | 0.44 | 1.88E-02 | 0.46 | -0.04 | -0.19 |
| DHRS9 | 0.44 | 1.48E-02 | 0.38 | 0.70 | -0.25 |
| EIF4G1 | 0.44 | 2.86E-02 | 0.30 | -0.33 | 0.29 |
| DHX9 // NPL | 0.43 | 3.62E-03 | 0.31 | 1.24 | -1.03 |
| HELLS | 0.43 | 3.01E-02 | -0.11 | -0.78 | -0.80 |
| PIGW | 0.43 | 4.49E-02 | 0.08 | 0.12 | -0.77 |
| CRYM | 0.43 | 3.31E-02 | 0.49 | 1.88 | -0.76 |
| CKS2 | 0.43 | 3.29E-02 | -0.15 | -0.75 | -0.51 |
| WDR48 | 0.43 | 2.57E-02 | 0.38 | 0.48 | -0.34 |
| UTP18 | 0.43 | 2.42E-03 | 0.39 | -0.18 | -0.12 |
| FUNDC1 | 0.43 | 2.50E-02 | 0.00 | 0.30 | -0.16 |
| RFK | 0.43 | 4.07E-02 | 0.21 | 0.33 | -0.07 |
| ABHD11 | 0.43 | 1.25E-02 | 0.51 | 0.37 | 0.11 |
| GPR37 | 0.42 | 1.44E-02 | -0.04 | 1.12 | -1.16 |
| VILL | 0.42 | 1.18E-02 | 0.56 | 0.16 | -1.12 |
| CNTN1 | 0.42 | 1.98E-02 | 0.23 | -0.51 | -0.93 |
| ATP1B3 | 0.42 | 1.75E-02 | -0.15 | 0.26 | -0.83 |
| CMAHP | 0.42 | 4.40E-02 | 0.93 | 2.05 | -0.96 |
| NAA15 | 0.42 | 3.60E-03 | 0.19 | -0.26 | -0.42 |
| CYCS | 0.42 | 8.80E-04 | -0.31 | 0.24 | -0.48 |
| PMS2P4 | 0.42 | 2.69E-02 | 0.06 | Not Detected | -0.36 |
| ALG11 // UTP14C | 0.42 | 2.96E-02 | 0.10 | 1.01 | -0.31 |
| ZNRF2 | 0.42 | 7.96E-03 | -0.16 | -0.25 | -0.15 |
| FAM199X | 0.42 | 3.38E-03 | -0.16 | 0.06 | -0.04 |
| PIGN | 0.42 | 3.60E-04 | 0.31 | 0.64 | -0.24 |
| CCAR2 | 0.42 | 2.65E-02 | 0.16 | 0.21 | -0.10 |
| AZI2 | 0.42 | 5.56E-03 | 0.06 | 0.37 | -0.12 |
| TBK1 | 0.42 | 1.80E-04 | 0.19 | 0.32 | -0.06 |
| N4BP2 | 0.42 | 1.61E-02 | 0.08 | 0.18 | -0.15 |
| LGALS9B | 0.42 | 3.74E-02 | 0.50 | 0.95 | -0.12 |
| DDX39A | 0.42 | 5.54E-03 | 0.08 | -1.07 | 0.03 |
| TRMT6 | 0.42 | 4.65E-02 | 0.12 | -0.48 | 0.12 |
| TIAL1 | 0.42 | 1.48E-02 | 0.03 | -0.28 | 0.23 |
| CEP295 | 0.42 | 3.45E-02 | 0.18 | -0.06 | 0.08 |
| HLA-DOB // TAP2 | 0.42 | 3.78E-02 | -0.39 | 0.28 | 0.10 |
| MUL1 | 0.42 | 2.07E-02 | 0.24 | 0.12 | 0.10 |
| MORC3 | 0.42 | 3.44E-02 | 0.48 | 0.40 | 0.08 |
| BPNT1 | 0.42 | 4.02E-02 | 0.24 | 0.16 | 0.30 |
| ARHGEF10L | 0.42 | 2.00E-02 | 0.15 | Not Detected | 0.19 |
| NASP | 0.41 | 3.34E-02 | 0.12 | -0.39 | -0.82 |
| STK17B | 0.41 | 1.40E-02 | 0.72 | 0.51 | -0.41 |
| DOPEY1 | 0.41 | 6.40E-03 | 0.53 | 0.50 | -0.34 |
| LLPH // TMBIM4 | 0.41 | 3.87E-02 | 0.12 | 0.79 | -0.41 |
| SMC2 | 0.41 | 1.27E-02 | 0.11 | -0.64 | -0.10 |
| MSH6 | 0.41 | 2.65E-02 | 0.04 | 0.01 | -0.19 |
| C10orf76 | 0.41 | 2.50E-02 | 0.16 | 0.07 | -0.08 |
| SETDB1 | 0.41 | 3.37E-02 | 0.18 | -0.12 | 0.08 |
| SNIP1 | 0.41 | 4.04E-02 | 0.59 | 0.26 | 0.29 |
| ALDH3A1 | 0.40 | 1.53E-02 | 1.12 | -1.14 | -3.01 |
| NDUFB1 | 0.40 | 3.36E-02 | 0.10 | 0.20 | -0.59 |
| ZNF19 // ZNF23 | 0.40 | 8.50E-03 | -0.11 | 0.14 | -0.26 |
| ACAD8 | 0.40 | 4.95E-02 | 0.19 | 0.42 | -0.26 |
| LRRC16A | 0.40 | 3.29E-02 | 0.29 | 0.41 | -0.58 |
| EREG | 0.40 | 3.04E-02 | 0.53 | -1.52 | -0.20 |
| PPCS | 0.40 | 2.29E-02 | 0.32 | 0.88 | -0.24 |
| ZCRB1 | 0.40 | 8.80E-03 | 0.28 | 0.40 | -0.06 |
| FIG4 | 0.40 | 6.10E-03 | 0.25 | 0.18 | -0.04 |
| RALGAPA2 | 0.40 | 4.53E-02 | 0.65 | 0.74 | -0.24 |
| EPS8L1 | 0.40 | 3.32E-02 | 0.60 | -0.50 | 0.24 |
| KCMF1 | 0.40 | 2.42E-02 | -0.29 | -0.01 | 0.06 |
| KIF21A | 0.40 | 4.42E-02 | 0.00 | 0.38 | 0.04 |
| SRSF5 | 0.40 | 2.50E-02 | 0.25 | 0.14 | 0.24 |
| POC5 | 0.40 | 4.70E-03 | 0.06 | 0.08 | 0.29 |
| CRNKL1 | 0.40 | 3.10E-02 | 0.28 | 0.32 | 0.06 |
| GCLC | 0.39 | 3.12E-03 | 0.46 | -0.32 | -0.93 |
| PLA2G4A | 0.39 | 3.81E-02 | 0.48 | 0.19 | -0.65 |
| XBP1 | 0.39 | 1.59E-02 | 0.45 | 1.06 | -0.59 |
| SCEL | 0.39 | 1.85E-02 | 0.06 | -0.71 | -0.55 |
| RPAP3 | 0.39 | 4.68E-02 | 0.45 | -0.30 | -0.34 |
| UFL1 | 0.39 | 2.63E-02 | 0.37 | 0.20 | -0.28 |
| GRHL2 | 0.39 | 5.00E-04 | 0.71 | 0.34 | -0.43 |
| C1orf21 | 0.39 | 4.83E-02 | 0.06 | 0.31 | -0.52 |
| DKC1 | 0.39 | 1.60E-03 | 0.04 | -0.80 | -0.24 |
| CCDC125 | 0.39 | 2.82E-02 | 0.29 | -0.20 | -0.07 |
| ALG1 // NAGPA-AS1 | 0.39 | 4.83E-02 | 0.39 | 0.06 | -0.14 |
| COIL | 0.39 | 3.29E-02 | 0.31 | -0.29 | 0.34 |
| MTRF1L | 0.39 | 3.12E-03 | 0.15 | -0.16 | 0.08 |
| G025368_T107514 | 0.39 | 7.46E-03 | Not Detected | Not Detected | Not Detected |
| FEN1 | 0.38 | 2.94E-02 | 0.37 | -0.68 | -0.61 |
| KRT15 | 0.38 | 3.61E-02 | 1.29 | 0.39 | -0.84 |
| CHKA | 0.38 | 4.13E-02 | 0.28 | 0.86 | -0.33 |
| PAK1IP1 | 0.38 | 1.85E-02 | -0.15 | -0.29 | -0.14 |
| PNO1 | 0.38 | 2.84E-02 | 0.36 | -0.49 | -0.04 |
| PSMD1 | 0.38 | 5.64E-03 | 0.08 | -0.28 | -0.03 |
| GTF2H2 | 0.38 | 6.36E-03 | -0.46 | -0.01 | -0.24 |
| PIGV | 0.38 | 3.05E-02 | 0.16 | 0.18 | -0.24 |
| SECISBP2 | 0.38 | 5.60E-03 | 0.14 | 0.21 | -0.23 |
| NRDE2 | 0.38 | 2.73E-02 | 0.28 | 0.26 | -0.10 |
| KLK5 | 0.38 | 2.46E-02 | 0.30 | -1.66 | 0.29 |
| HSP90AB1 | 0.38 | 4.52E-02 | -0.11 | -0.44 | 0.19 |
| ACTL6A | 0.38 | 5.00E-02 | -0.06 | -0.20 | 0.24 |
| GALNT10 | 0.38 | 3.96E-03 | 0.50 | 0.00 | 0.73 |
| ZAP70 | 0.38 | 1.31E-02 | Not Detected | Not Detected | Not Detected |
| RFC4 | 0.37 | 4.18E-02 | 0.11 | -0.50 | -0.50 |
| TUBD1 | 0.37 | 1.48E-03 | -0.12 | 0.31 | -0.28 |
| DNAJA4 | 0.37 | 7.08E-03 | 0.26 | 0.39 | -0.38 |
| ATP7A | 0.37 | 3.71E-02 | 0.29 | 0.69 | -0.33 |
| C9orf78 | 0.37 | 4.20E-02 | 0.25 | -0.08 | -0.03 |
| AP3B1 | 0.37 | 3.72E-02 | 0.11 | 0.38 | -0.25 |
| COMMD10 | 0.37 | 3.24E-02 | 0.01 | 0.06 | -0.16 |
| UFC1 | 0.37 | 2.74E-02 | -0.01 | -0.39 | 0.24 |
| TASP1 | 0.37 | 1.91E-02 | 0.24 | -0.41 | 0.37 |
| FGFR1OP2 | 0.37 | 4.50E-02 | -0.10 | -0.04 | 0.25 |
| DUSP4 | 0.37 | 4.28E-02 | 0.77 | -0.16 | 0.56 |
| AGA | 0.37 | 2.01E-02 | -0.33 | 0.23 | 0.34 |
| STX18 | 0.37 | 3.03E-02 | 0.14 | 0.23 | 0.06 |
| TRIM24 | 0.37 | 3.47E-02 | 0.57 | 0.30 | 0.10 |
| TCEAL9 | 0.37 | 4.22E-02 | 0.45 | 0.16 | 0.30 |
| OR2A4 | 0.37 | 3.52E-02 | Not Detected | Not Detected | 0.77 |
| SASS6 | 0.36 | 1.19E-02 | 0.32 | -0.44 | -0.58 |
| MAN2A2 | 0.36 | 2.05E-02 | 0.28 | -0.30 | -0.42 |
| GALNT8 // NDUFA9 | 0.36 | 4.32E-02 | -0.03 | -0.06 | -0.32 |
| UHMK1 | 0.36 | 3.62E-02 | 0.40 | 0.62 | -0.51 |
| ALPK1 | 0.36 | 1.70E-02 | 0.41 | 0.88 | -0.55 |
| ESF1 | 0.36 | 1.30E-02 | 0.06 | 0.11 | -0.31 |
| GSN | 0.36 | 1.03E-02 | 0.77 | 1.39 | -0.38 |
| SECISBP2L | 0.36 | 2.10E-02 | 0.37 | 1.14 | -0.21 |
| C12orf4 | 0.36 | 8.36E-03 | 0.34 | 0.10 | -0.01 |
| DCUN1D3 // ERI2 | 0.36 | 1.73E-02 | 0.26 | 0.03 | -0.18 |
| PXYLP1 | 0.36 | 2.27E-02 | 0.18 | 0.77 | -0.16 |
| SLC6A14 | 0.36 | 4.14E-02 | 0.04 | 0.40 | -0.20 |
| MRPS7 | 0.36 | 2.45E-02 | -0.07 | -0.39 | 0.08 |

| | KD Normal (Mean) | | KD COPD (Mean) | Diff Normal (Mean) | TGFβ Normal (Mean) |
|---|---|---|---|---|---|
| Gene Symbol | Fold-Change (Log2) | P-value | Fold-Change (Log2) | Fold-Change (Log2) | Fold-Change (Log2) |
| PWP1 | 0.36 | 5.12E-03 | 0.04 | -0.37 | 0.19 |
| SLC12A2 | 0.36 | 4.81E-02 | 0.08 | 0.24 | 0.12 |
| FOXC1 | 0.36 | 1.09E-02 | 0.56 | 0.14 | 0.18 |
| ALDH3A2 | 0.34 | 4.89E-02 | 0.20 | 0.26 | -1.03 |
| HIST2H4A // HIST2H4B | 0.34 | 1.02E-03 | 0.49 | -0.63 | -0.44 |
| EI24 | 0.34 | 4.57E-02 | -0.10 | -0.08 | -0.34 |
| G064937_T279680 | 0.34 | 4.06E-02 | 0.04 | 0.57 | -0.26 |
| U2SURP | 0.34 | 2.78E-03 | 0.10 | 0.03 | -0.28 |
| RMND5A | 0.34 | 3.82E-02 | 0.33 | 0.68 | -0.41 |
| TSPAN8 | 0.34 | 1.82E-02 | 0.64 | 1.05 | -0.38 |
| PHLPP1 | 0.34 | 7.48E-03 | 0.37 | 0.03 | -0.37 |
| DEFB1 | 0.34 | 1.13E-02 | 0.32 | 0.98 | -0.42 |
| NOP58 | 0.34 | 1.23E-02 | -0.11 | -0.51 | -0.16 |
| NUP155 | 0.34 | 8.00E-04 | 0.01 | -0.57 | -0.25 |
| APTX | 0.34 | 9.70E-03 | 0.41 | -0.39 | -0.11 |
| EED | 0.34 | 3.92E-03 | 0.28 | -0.08 | -0.14 |
| G081924_T347654 | 0.34 | 2.56E-03 | -0.26 | 0.80 | -0.14 |
| AGPS | 0.34 | 3.14E-02 | -0.19 | 0.03 | -0.08 |
| NR6A1 | 0.34 | 1.43E-02 | -0.18 | 0.00 | -0.04 |
| ERLIN2 | 0.34 | 1.62E-02 | 0.25 | 0.43 | -0.18 |
| ZNF24 | 0.34 | 4.92E-02 | 0.38 | 0.28 | -0.11 |
| TAOK3 | 0.34 | 4.18E-02 | 0.46 | 0.16 | -0.03 |
| RTN3 | 0.34 | 4.30E-02 | 0.40 | 0.15 | -0.19 |
| SERPINB2 | 0.34 | 4.90E-02 | 0.14 | -1.71 | 0.57 |
| MRPL1 | 0.34 | 7.10E-03 | -0.19 | -0.53 | 0.10 |
| CDK6 | 0.34 | 7.78E-03 | 0.75 | -0.38 | 0.38 |
| RNF122 | 0.34 | 4.62E-03 | -0.06 | Not Detected | 0.78 |
| NAA60 | 0.34 | 4.05E-02 | 0.15 | 0.04 | 0.16 |
| SLC25A26 | 0.34 | 4.32E-02 | 0.01 | 0.01 | 0.18 |
| MUC1 | 0.33 | 1.47E-02 | 0.33 | 0.99 | -0.94 |
| CNBP | 0.33 | 2.74E-02 | 0.07 | -0.07 | -0.31 |
| SPRYD4 | 0.33 | 3.58E-02 | 0.12 | -0.06 | -0.53 |
| NPAT | 0.33 | 4.93E-02 | 0.04 | 0.33 | -0.31 |
| LINC01881 | 0.33 | 4.79E-02 | 0.28 | 0.66 | -0.41 |
| YTHDC2 | 0.33 | 2.87E-02 | 0.28 | 0.14 | -0.40 |
| CUL3 | 0.33 | 4.52E-02 | 0.01 | 0.41 | -0.39 |
| PPP1CC | 0.33 | 1.43E-02 | 0.01 | 0.06 | -0.32 |
| PCCB | 0.33 | 4.07E-02 | 0.25 | 0.00 | -0.20 |
| EDARADD | 0.33 | 4.94E-02 | -0.07 | 0.34 | -0.25 |
| RUFY3 | 0.33 | 3.89E-02 | 0.18 | 0.39 | -0.12 |
| GPBP1 | 0.33 | 4.42E-02 | -0.20 | 0.14 | 0.07 |
| PTP4A2 | 0.33 | 3.08E-02 | 0.01 | 0.41 | 0.08 |
| POLR2A | 0.33 | 9.80E-03 | 0.44 | 0.11 | 0.38 |
| ATP6V0E2 | 0.33 | 2.52E-02 | 0.82 | Not Detected | Not Detected |
| CDKN3 | 0.32 | 3.37E-02 | -0.04 | -0.69 | -1.93 |
| AKR1B10 | 0.32 | 2.30E-02 | 0.48 | -0.63 | -1.62 |
| HJURP | 0.32 | 1.72E-02 | 0.10 | -0.75 | -0.94 |
| KDELC2 | 0.32 | 4.12E-02 | -0.14 | -0.18 | -0.74 |
| CLN6 | 0.32 | 1.97E-02 | 0.06 | -0.16 | -0.71 |
| REEP3 | 0.32 | 4.33E-02 | 0.37 | 0.76 | -0.31 |
| TSPAN3 | 0.32 | 2.74E-03 | 0.06 | 0.42 | -0.40 |
| GGCX | 0.32 | 3.90E-02 | 0.40 | 0.14 | -0.32 |
| PSMC1 | 0.32 | 5.46E-03 | -0.14 | -0.37 | -0.15 |
| VPS29 | 0.32 | 9.40E-04 | -0.42 | -0.21 | -0.24 |
| STK17A | 0.32 | 1.22E-02 | -0.08 | -0.21 | -0.03 |
| SLC44A3 | 0.32 | 4.35E-02 | -0.15 | 0.32 | -0.14 |
| USP16 | 0.32 | 1.82E-02 | -0.06 | 0.12 | -0.25 |
| PNPLA8 | 0.32 | 3.93E-02 | 0.15 | 0.40 | -0.21 |
| VAMP7 | 0.32 | 2.24E-02 | 0.11 | 0.07 | -0.01 |
| RALBP1 | 0.32 | 4.23E-02 | 0.44 | 0.12 | -0.10 |
| CTAGE9 | 0.32 | 2.02E-02 | 0.48 | 0.55 | -0.01 |
| HIST2H2AB // HIST2H2E | 0.32 | 2.86E-02 | 0.01 | 0.15 | -0.21 |
| TMEM156 | 0.32 | 4.63E-02 | -0.18 | -1.14 | 0.43 |
| SMIM12 | 0.32 | 2.00E-04 | -0.18 | -0.21 | 0.19 |
| UTP6 | 0.32 | 1.12E-02 | 0.37 | -0.19 | 0.29 |
| PMS2P3 | 0.32 | 3.73E-02 | -0.21 | Not Detected | 0.36 |
| EIF4E2 | 0.32 | 1.27E-02 | 0.69 | 0.04 | 0.03 |
| BCAP29 | 0.32 | 3.62E-02 | 0.18 | 0.37 | 0.12 |
| OSBPL8 | 0.32 | 2.30E-02 | 0.25 | 0.23 | 0.10 |
| CNST | 0.32 | 1.80E-02 | 0.16 | 0.40 | 0.08 |
| STK38 | 0.32 | 1.06E-02 | 0.25 | 0.56 | 0.20 |
| CHST9 | 0.31 | 3.69E-02 | 1.34 | 1.37 | -1.34 |
| PLRG1 | 0.31 | 1.96E-03 | 0.30 | 0.68 | -0.64 |
| TIMM21 | 0.31 | 2.50E-02 | 0.19 | -0.15 | -0.55 |
| TMBIM6 | 0.31 | 1.00E-02 | 0.28 | 0.18 | -0.36 |
| SLC25A4 | 0.31 | 4.80E-02 | 0.42 | 0.23 | -0.29 |
| AGL | 0.31 | 2.66E-02 | 0.18 | 0.16 | -0.44 |
| UBR7 | 0.31 | 4.22E-02 | 0.56 | 0.34 | -0.43 |
| TUBGCP5 | 0.31 | 6.28E-03 | 0.08 | -0.32 | -0.04 |
| NCCRP1 | 0.31 | 2.59E-02 | 0.56 | -0.42 | -0.23 |
| RBM25 | 0.31 | 2.49E-02 | 0.06 | -0.15 | -0.12 |
| TOM1L2 | 0.31 | 3.52E-02 | 0.34 | -0.06 | -0.25 |
| THUMPD3 | 0.31 | 4.15E-02 | 0.24 | -0.08 | -0.18 |
| NMD3 | 0.31 | 3.48E-02 | 0.14 | -0.15 | -0.10 |
| SLMAP | 0.31 | 3.83E-02 | 0.07 | 0.31 | -0.07 |
| CIRBP | 0.31 | 1.85E-02 | 0.77 | 0.77 | -0.25 |
| ARMCX6 | 0.31 | 4.85E-02 | -0.06 | -0.39 | 0.46 |
| MTG2 | 0.31 | 1.65E-02 | 0.38 | -0.54 | 0.24 |
| LARS | 0.31 | 3.05E-02 | -0.32 | -0.11 | 0.01 |
| INSIG2 | 0.31 | 1.69E-02 | 0.04 | 0.12 | 0.08 |
| OCRL | 0.31 | 1.24E-02 | 0.20 | 0.23 | 0.01 |
| PRIM1 | 0.30 | 3.66E-03 | -0.10 | -0.73 | -1.08 |
| DKK1 | 0.30 | 1.26E-02 | 0.32 | -1.35 | -0.82 |
| PIR // VEGFD | 0.30 | 4.13E-02 | 0.10 | -0.63 | -0.71 |
| TNFRSF21 | 0.30 | 2.66E-02 | 0.45 | 0.71 | -0.86 |
| EBP | 0.30 | 3.85E-02 | -0.32 | -0.43 | -0.46 |
| ANAPC11 | 0.30 | 1.64E-02 | -0.16 | -0.20 | -0.26 |
| PIGU | 0.30 | 1.29E-02 | 0.01 | -0.10 | -0.36 |
| IFT57 | 0.30 | 4.35E-02 | 0.28 | 0.10 | -0.33 |
| EPHX1 | 0.30 | 8.54E-03 | 0.90 | 1.06 | -0.55 |
| PFDN2 | 0.30 | 2.79E-02 | 0.18 | -0.24 | -0.06 |
| TMEM9B | 0.30 | 3.51E-02 | 0.26 | 0.39 | -0.14 |
| PARP9 | 0.30 | 1.66E-02 | 0.28 | 0.36 | -0.25 |
| ACER3 | 0.30 | 4.16E-02 | -0.14 | -0.54 | 0.04 |
| MRPL18 | 0.30 | 7.80E-04 | 0.14 | -0.28 | 0.11 |
| AKT1 | 0.30 | 2.79E-02 | 0.07 | -0.14 | 0.33 |
| LARS2 | 0.30 | 1.32E-02 | 0.29 | -0.23 | 0.29 |
| ADO | 0.30 | 4.84E-02 | -0.14 | 0.04 | 0.34 |
| RGS12 | 0.30 | 4.62E-02 | -0.03 | 0.10 | 0.30 |
| SNX14 | 0.30 | 1.00E-04 | 0.06 | 0.42 | 0.03 |
| TMEM114 | 0.30 | 2.64E-02 | 0.36 | Not Detected | Not Detected |
| PLEKHS1 | 0.29 | 3.63E-02 | 0.53 | 2.17 | -1.52 |
| C19orf48 | 0.29 | 3.23E-02 | 0.08 | -0.60 | -0.65 |
| RPAIN | 0.29 | 4.03E-02 | 0.01 | 0.21 | -0.60 |
| RNASEH2A | 0.29 | 1.09E-02 | 0.04 | -0.66 | -0.39 |
| NUP160 | 0.29 | 2.39E-02 | 0.19 | -0.06 | -0.39 |
| GSS | 0.29 | 3.15E-02 | 0.00 | 0.21 | -0.48 |
| TMEM50B | 0.29 | 2.88E-03 | 0.12 | 0.59 | -0.44 |
| ZNF638 | 0.29 | 1.83E-02 | 0.03 | 0.50 | -0.32 |
| MIPEP | 0.29 | 7.76E-03 | 0.14 | 0.19 | -0.51 |
| TPD52L1 | 0.29 | 1.05E-02 | 0.04 | -0.12 | -0.04 |
| SVIP | 0.29 | 9.74E-03 | 0.25 | -0.19 | -0.15 |
| KPNB1 | 0.29 | 1.44E-02 | 0.10 | -0.10 | -0.08 |
| TMEM38B | 0.29 | 4.31E-02 | 0.42 | -0.15 | -0.21 |
| NSA2 | 0.29 | 1.28E-02 | -0.10 | 0.00 | -0.20 |
| TM9SF3 | 0.29 | 4.74E-02 | 0.01 | 0.30 | -0.11 |
| LMBR1 | 0.29 | 1.54E-02 | 0.01 | 0.42 | -0.21 |
| CASP3 | 0.29 | 1.97E-02 | 0.11 | 0.40 | -0.16 |
| WIPI1 | 0.29 | 1.55E-02 | 0.07 | 0.34 | -0.14 |
| GNL2 | 0.29 | 1.70E-02 | -0.06 | -0.29 | 0.10 |
| CTC1 | 0.29 | 8.90E-03 | 0.19 | -0.53 | Not Detected |
| G084587_T358508 | 0.29 | 3.54E-02 | 0.61 | -0.23 | Not Detected |
| SIL1 | 0.29 | 2.05E-02 | 0.11 | -0.01 | 0.19 |
| FBXO30 | 0.29 | 6.20E-04 | 0.08 | 0.07 | 0.31 |
| MOCS3 | 0.29 | 3.15E-02 | 0.14 | 0.03 | 0.08 |
| AURKAPS1 // RAB3GAF | 0.29 | 1.18E-02 | 0.10 | 0.29 | 0.14 |
| IFT20 | 0.29 | 3.13E-02 | 0.15 | 0.51 | 0.01 |
| SLC39A3 | 0.29 | 1.07E-02 | 0.59 | 0.01 | 0.29 |
| CDCA7L | 0.28 | 1.14E-02 | -0.06 | -0.82 | -0.36 |
| ERGIC2 | 0.28 | 1.54E-02 | 0.23 | 0.16 | -0.26 |
| SF3A3 | 0.28 | 8.20E-04 | 0.20 | 0.21 | -0.31 |
| UBE4A | 0.28 | 1.85E-02 | 0.10 | 0.41 | -0.51 |
| USP28 | 0.28 | 4.88E-02 | 0.34 | 0.38 | -0.58 |
| ADGRL1 | 0.28 | 4.15E-02 | 0.60 | 1.30 | -0.28 |
| ALDH1L1 | 0.28 | 2.02E-02 | 0.77 | 1.16 | -0.26 |
| GEN1 | 0.28 | 1.08E-03 | 0.01 | -0.40 | -0.16 |
| PNPT1 | 0.28 | 9.06E-03 | 0.23 | -0.45 | -0.20 |
| GATC | 0.28 | 6.52E-03 | 0.08 | -0.19 | -0.10 |
| CRAMP1 // HN1L | 0.28 | 2.13E-02 | 0.37 | -0.01 | -0.11 |
| SLC2A10 | 0.28 | 4.88E-02 | 0.48 | 0.71 | -0.06 |
| FNBP4 | 0.28 | 3.70E-02 | 0.15 | 0.10 | -0.18 |
| STYX | 0.28 | 2.08E-02 | 0.18 | -0.08 | 0.08 |
| CDK12 | 0.28 | 2.10E-02 | 0.11 | -0.04 | 0.20 |
| SAMD12-AS1 | 0.28 | 1.96E-02 | 0.06 | -0.06 | 0.25 |
| NBPF14 | 0.28 | 1.28E-02 | 0.04 | 0.16 | 0.65 |
| ABHD13 | 0.28 | 4.35E-02 | 0.03 | 0.21 | 0.33 |
| DIS3L | 0.26 | 3.70E-02 | 0.34 | 0.14 | -0.70 |
| MTHFD1 | 0.26 | 4.68E-02 | 0.24 | -0.18 | -0.57 |
| SLC35C1 | 0.26 | 3.38E-02 | 0.29 | 0.67 | -0.51 |
| TTC22 | 0.26 | 2.28E-02 | 0.08 | 0.01 | -0.26 |
| CCT7 | 0.26 | 4.36E-02 | -0.10 | -0.41 | -0.19 |
| ME2 | 0.26 | 4.59E-02 | -0.08 | -0.25 | -0.07 |
| NSL1 | 0.26 | 1.40E-02 | -0.04 | 0.18 | -0.14 |
| FBXL4 | 0.26 | 1.05E-02 | 0.14 | 0.33 | -0.03 |
| TBRG1 | 0.26 | 3.48E-03 | 0.10 | 0.08 | -0.19 |
| TJP3 | 0.26 | 2.25E-02 | 0.52 | 0.51 | -0.19 |
| PPRC1 | 0.26 | 2.68E-02 | 0.12 | -0.54 | 0.03 |
| L3MBTL3 | 0.26 | 2.65E-02 | 0.21 | -0.04 | 0.74 |
| ACTR10 | 0.26 | 1.26E-02 | 0.06 | 0.04 | 0.14 |
| BEX3 | 0.26 | 2.31E-02 | 0.14 | 0.00 | 0.00 |

Genes statistically significant upregulated by Slug knockdown in normal bronchial basal/progenitor cells:

Table shows the response of the genes to differentiation and TGFβ and fold-change for Slug knockdown in COPD cells. Genes were selected for a log2 (fold-Change) ≥0.26. Statistical significance is at p-val < 5.00E-02

Figure S1

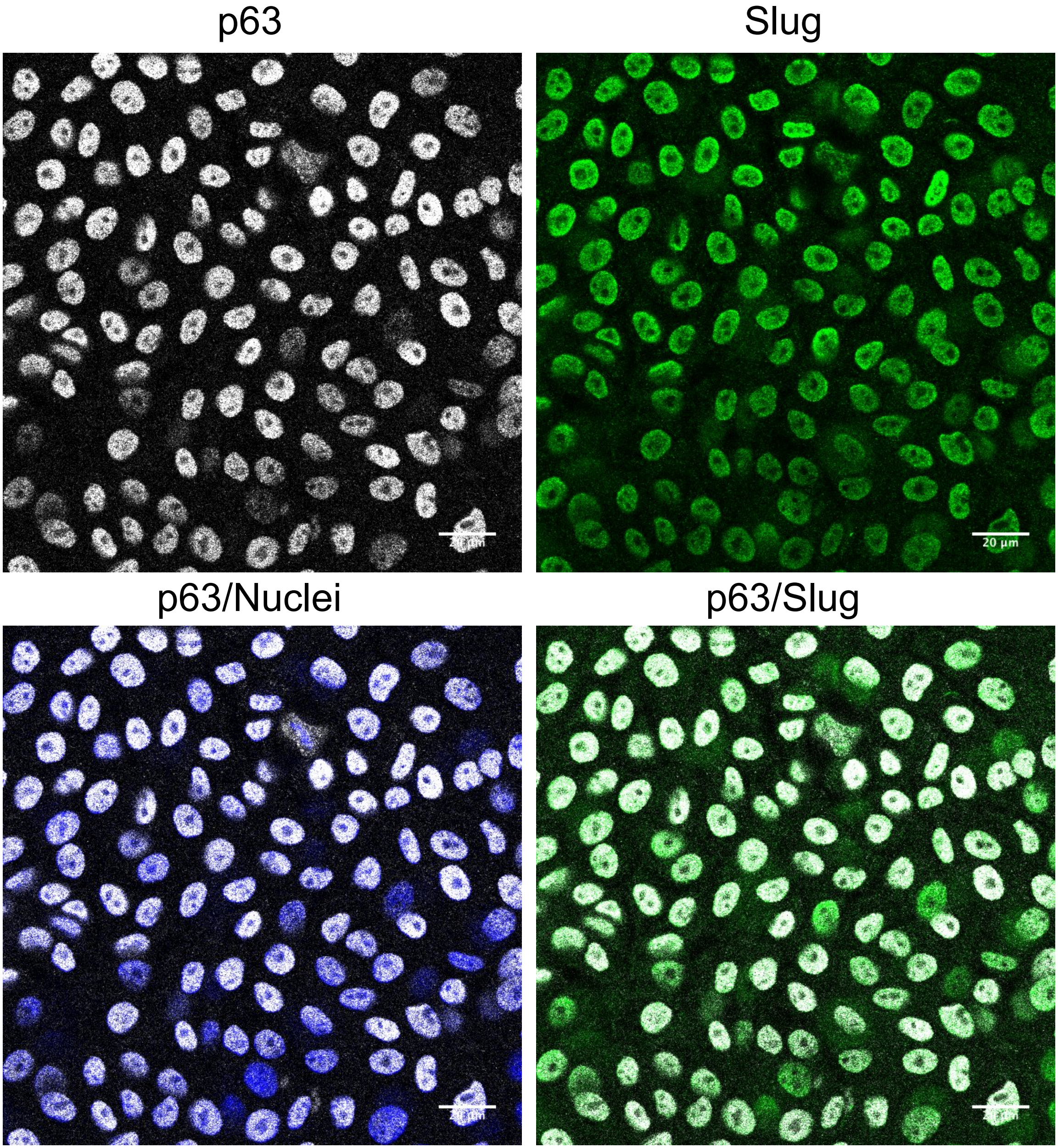

Figure S2

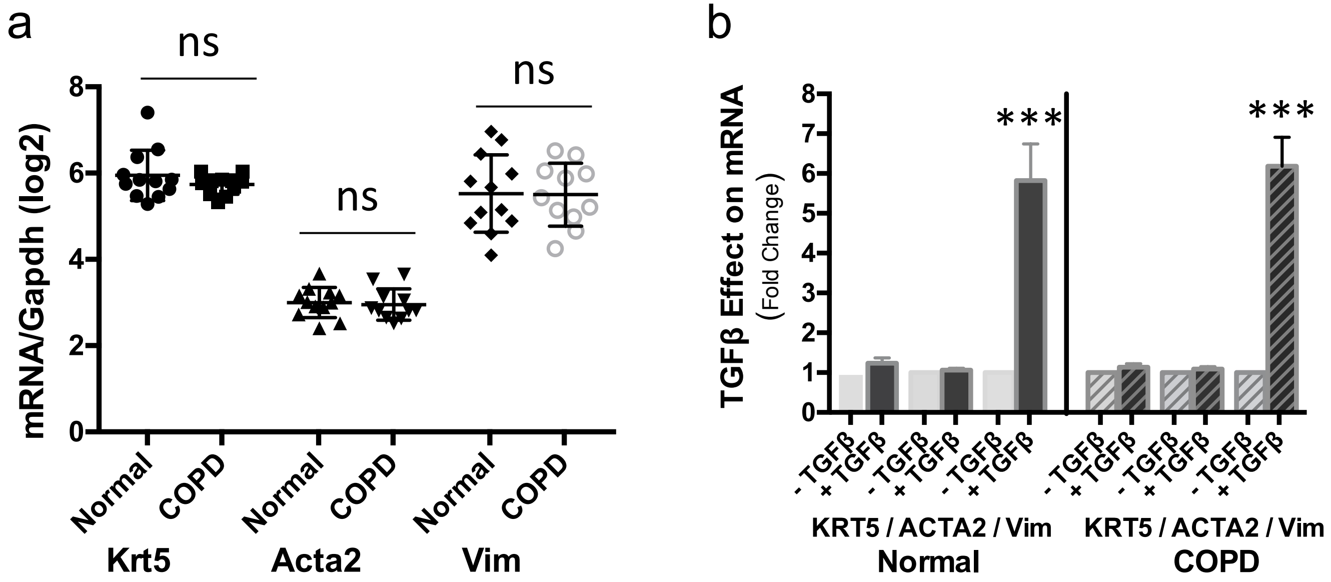
a
ns
ns
ns
mRNA/Gapdh (log2)
8
6
4
2
0
Normal
COPD
Normal
COPD
Normal
COPD
Krt5
Acta2
Vim
b
TGFβ Effect on mRNA
(Fold Change)
8
7
6
5
4
3
2
1
0
***
***
- TGFβ
+ TGFβ
- TGFβ
+ TGFβ
- TGFβ
+ TGFβ
- TGFβ
+ TGFβ
- TGFβ
+ TGFβ
- TGFβ
+ TGFβ
KRT5 / ACTA2 / Vim
KRT5 / ACTA2 / Vim
Normal
COPD

Figure S3

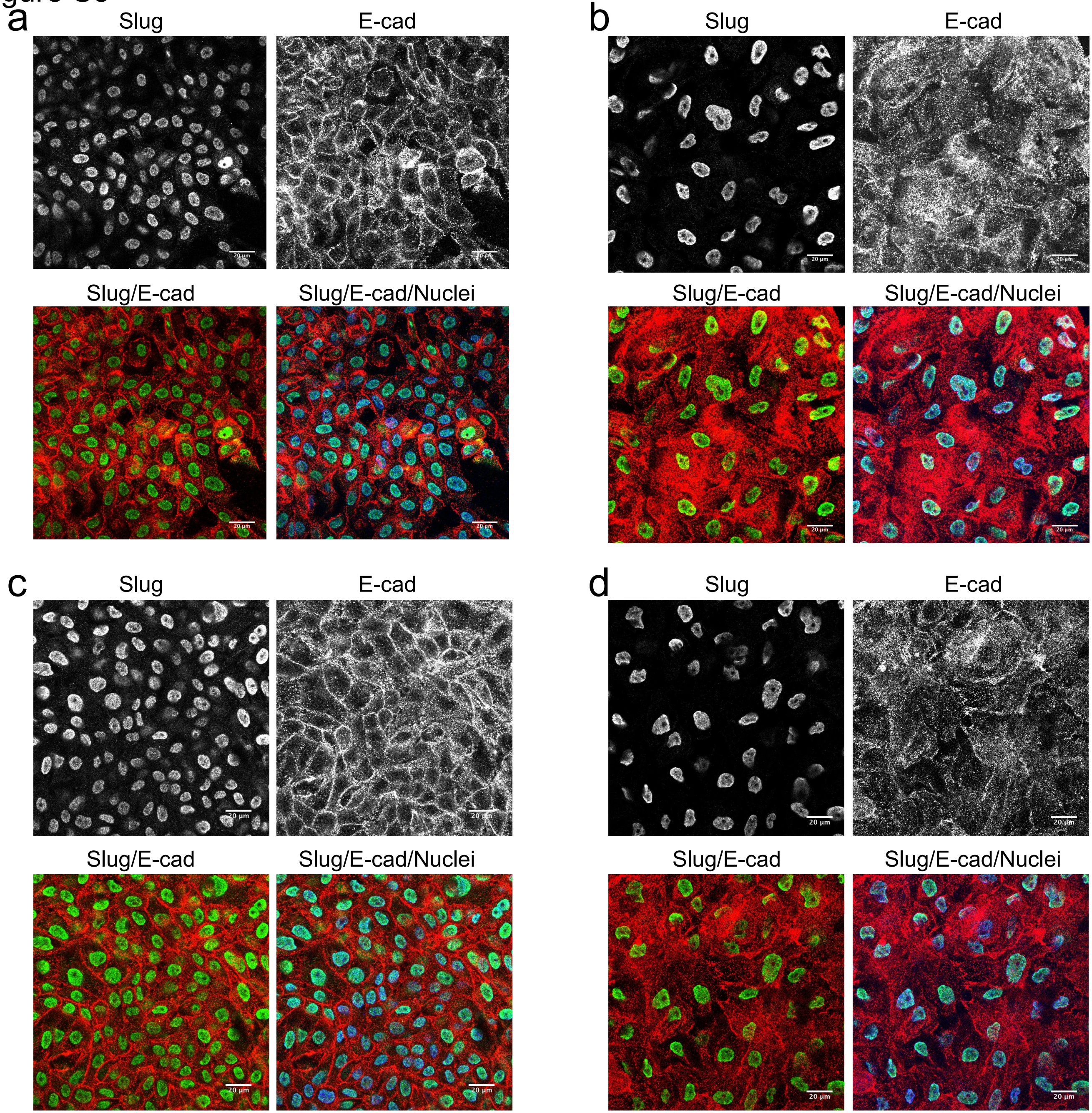
a
Slug
E-cad
Slug/E-cad
Slug/E-cad/Nuclei
b
Slug
E-cad
Slug/E-cad
Slug/E-cad/Nuclei
c
Slug
E-cad
Slug/E-cad
Slug/E-cad/Nuclei
d
Slug
E-cad
Slug/E-cad
Slug/E-cad/Nuclei
20 µm

Figure S4

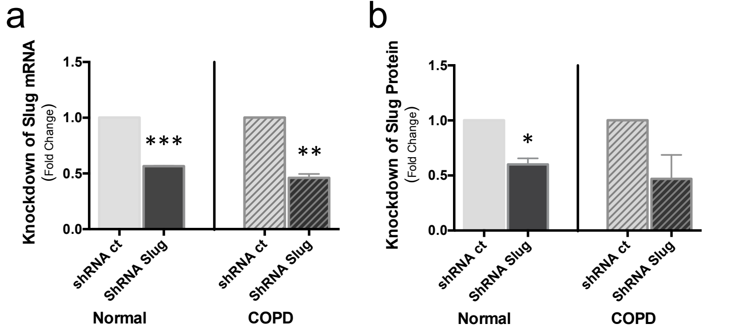
a
Knockdown of Slug mRNA
(Fold Change)
1.5
1.0
0.5
0.0
***
**
shRNA ct
ShRNA Slug
shRNA ct
ShRNA Slug
Normal
COPD
b
Knockdown of Slug Protein
(Fold Change)
1.5
1.0
0.5
0.0
*
shRNA ct
ShRNA Slug
shRNA ct
ShRNA Slug
Normal
COPD

### Figure S5

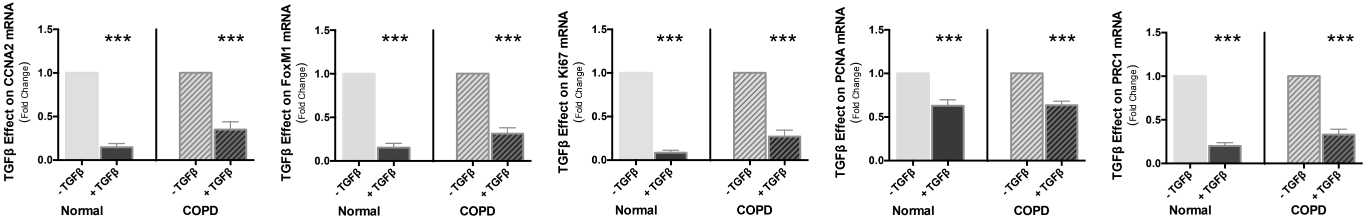

TGFβ Effect on CCNA2 mRNA
(Fold Change)
TGFβ Effect on FoxM1 mRNA
(Fold Change)
TGFβ Effect on Ki67 mRNA
(Fold Change)
TGFβ Effect on PCNA mRNA
(Fold Change)
TGFβ Effect on PRC1 mRNA
(Fold Change)
0.0
0.5
1.0
1.5
***
- TGFβ
+ TGFβ
Normal
COPD

### Supplementary Figure Legends

**Fig. S1 Bronchial epithelial progenitors from COPD express Slug in their nuclei.**

Primary bronchial epithelial basal cells from COPD were grown on filters and analyzed undifferentiated at confluence by fluorescent immunocytochemistry. Cells were fixed and labeled simultaneously with progenitor cell marker p63 (white) and Slug (green) antibodies and with Hoechst as a marker of nuclei (blue). Bars are 20 μm.

**Fig. S2 Bronchial epithelial progenitors co-express epithelial and mesenchymal markers.**

Primary bronchial epithelial basal cells, normal and COPD, were grown on filters at confluence and either analyzed undifferentiated (**a**) or changed to ALI culture to induce differentiation, without TGF-β or in presence of 1ng/ml of TGF-β and analyzed at day 6 of ALI culture for mRNA expression (**b**). RNA were extracted from normal and COPD cells and analyzed by RT-qPCR to determine the mRNA levels of KRT5 gene, an epithelial cytoskeletal marker and ACTA 2 and Vim genes, mesenchymal cytoskeletal markers.. GAPDH was used to normalize cDNA amounts between samples and results were calculated as a ratio on GAPDH. Data are for $n \geq 11$ and experiments were done at least in duplicate. (**a**) Results shown are log2 (ratio on GAPDH) and are presented as a scatter plot with the mean ±SD. ns: non significant. (**b**) Results are presented as the fold-change induced by TGF-β on mRNA expression with mean ±SEM, and compare normal and COPD cells. Statistical significance is at P value < 1.00E-03 *** as indicated, ns is non significant.

**Fig. S3 Bronchial epithelial progenitors express Slug in their nuclei and co-express E-cadherin with or without TGF-β.** Normal (**a, b**) or COPD (**c, d**) primary bronchial epithelial basal cells were grown on filters and at confluence changed to ALI culture to induce differentiation, without TGF-β (**a, c**) or in presence of 1 ng/ml of TGF-β (**b, d**). Cells were analyzed at day 6 of ALI culture by fluorescent immunocytochemistry. Cells were fixed and labeled simultaneously with Slug (green) and E-cad (red) antibodies and with Hoechst as a marker of nuclei (blue). Bars are 20 μm.

**Fig. S4 Effect of Slug knockdown on Slug mRNA and protein levels in normal and COPD bronchial epithelial progenitors**

Normal and COPD bronchial epithelial basal/progenitor cells were transduced in transwell inserts with shRNA lentiviral particles corresponding to either a SNAI2/Slug specific sequence or control non-targeting sequences. At day 4 post-transduction, cells were changed to ALI conditions and at day 6 post-transduction cells were analyzed for mRNA or for protein expression. RNA or proteins lysates were prepared from normal and COPD cells and analyzed respectively by RT-qPCR (**a**) and by Western blot (**b**). For RT-qPCR analysis, GAPDH was used to normalize cDNA amounts between samples and results were calculated as a ratio on GAPDH. Results are presented as the fold-change induced by shRNA with SNAI2/Slug specific sequence on Slug mRNA expression (**a**) or Slug protein expression (**b**) with mean ±SEM. Data are for n $\geq$ 4. Statistical significance is at P value < 5.00E-02 *, < 1.00E-02 ** or < 1.00E-03 *** as indicated.

**Fig. S5 Comparison of TGF-β effect on the expression of proliferation-related genes between normal and COPD bronchial epithelial progenitors.**

Primary bronchial epithelial basal cells, normal and COPD, were grown on filters and analyzed undifferentiated at confluence or grown on filters and at confluence changed to ALI culture to induce differentiation, without TGF-β or in presence of 1 ng/ml of TGF-β. Cells were analyzed at day 6 of ALI culture for mRNA expression. RNA were prepared from normal and COPD cells and analyzed respectively by RT-qPCR. GAPDH was used to normalize cDNA amounts between samples and results were calculated as a ratio on GAPDH. Data shown are for n $\geq$ 11. Results are presented as the fold-change induced by TGF-β on mRNA expression with mean ±SEM. Statistical significance is at P value < 1.00E-03 *** as indicated.

#### S1 Appendix: Characteristics of the study subjects

| | non- COPD | | COPD | |
|---|---|---|---|---|
| | Non-smoker | Smoker | Moderate | Severe |
| **Subjects n** | 6 | 6 | 6 | 6 |
| **Female / male n** | 3/3 | 2/4 | 3/3 | 3/3 |
| **Age years** | 55 ±14.7 | 63 ±13.6 | 59 ±8.9 | 57 ±4.8 |
| **Smoking Status** | NA | Active | Active | Ex-Smoker |
| **Pack/year** | NA | 48 ±32.5 | 56 ±19.7 | 46 ±26.5 |
| **COPD Stage / FEV1%predicted** | NA | NA | GOLD2/74 ±5,7 | GOLD3-4 /28 ±11.4 |

Data are presented as Mean ±SD. Pack-year=1 year smoking 20 cigarettes per day. COPD, chronic obstructive pulmonary disease; FEV1, forced expiratory volume in 1 s;

#### S2 Appendix: Sequences of primers for PCR and shRNA

| Gene | NCBI | Forward primer | Reverse primer |
|---|---|---|---|
| CCNA2 | NM_001237 | 5'-TGT GGG CAC TGC TGC TAT GC -3' | 5'-GTG TCT CTG GTG GGT TGA GG -3' |
| FOXM1 | NM_202002 | 5'-AGC AAG CGA GTC CGC ATT GC-3' | 5'-GGA CCT AAG CCC ACT GTA GG-3' |
| VIM | NM_003380 | 5'-GTC CCT CAC CTG TGA AGT GG-3' | 5'-TTC CCT CAG GTT CAG GGA GG-3' |
| ACTA2 | NM_001141945 | 5'-AGA TCC TGA CTG AGC GTG GC -3' | 5'-TGC ATT CGG TCG GCA ATG CC -3' |
| CDH1 | NM_004360 | 5' GAG AGC GGT GGT CAA AGA GC 3' | 5' GAG GAG TTC AGG GAG CTC AG 3' |
| CDH2 | NM_001792 | 5'-TGC CAG TGT GAC TCC AAC GG-3' | 5'-TT CGT CGG ATT CCC ACA GGC-3' |
| GAPDH | NM_002046.6 | 5' TGT CAG TGG TGG ACC TGA CC 3' | 5' ACT CCT TGG AGG CCA TGT GG 3' |
| KRT5 | NM_000424.3 | 5'-CAA GGA TGC CAG GAA CAA GC -3' | 5'-CT GCT GGA GTA GTA GCT TCC -3' |
| MKi67 | NM_002417 | 5'-AGG GAA TAT CCC TGC GCT CC -3' | 5'-TCT CCT CTG CCA CCT TAG GC -3' |
| PCNA | NM_002592 | 5'-CCG AAG ATA ACG CGG ATA CC -3' | 5'-GTT GAA GAG AGT GGA GTG GC -3' |
| PRC1 | NM_003981 | 5'-GAC TGG CTC CCA ATA CAC CG -3' | 5'-GGA ACT GTC AGA GAG GGA CG -3' |
| SNAI1 | NM_005985 | 5'-AGG ACA GTG GGA AAG GCT CC-3' | 5'-ACA GGA GAA GGG CTT CTC GC-3' |
| SNAI2 | NM_003068 | 5'- AGC AGC TGC ACT GCG ATG CC -3' | 5' ACA CAG CAG CCA GAT TCC TC 3' |
| TWIST1 | NM_000474 | 5'-TCC GCA GTC TTA CGA GGA GC -3' | 5'-GTG AGC CAC ATA GCT GCA GC -3' |
| ZEB1 | NM_001128128 | 5'-TCA ACT ACG GTC AGC CCT GC -3' | 5'-GTG CTG TCA CGT TCT TCC GC -3' |
| **Gene** | **NCBI** | **Reference** | **Sequence** |
| shRNA SNAI2 | NM_003068 | TRCN0000284362: | CCGGGAGTGACGCAATCAATGTTTACTCGAGTAAACATYGATTGCGTCACTCTTTTTG |

**S3 Appendix: References for antibodies used in immunochemistry and RPPA**

| **Antibody** | **Supplier (Reference)** |
|---|---|
| **Antibodies for ICC** | |
| Alexa Fluor 488 | InVitrogen |
| Alexa Fluor 555 | InVitrogen |
| Alexa Fluor 647 | InVitrogen |
| E-Cadherin | CST (3195) |
| Slug/Snail2 | CST (9585) |
| P63 | DBS (4A4) |
| **Antibodies for RPPA** | |
| E-cadherin | BD (610181) |
| Slug | CST 9585) |
| Vimentin | CST (5741) |
| Ki67 | DAKO (M7240) |
| Smad3 | Epitomics (1735-1) |

CST: CST: Cell Signaling Technology; DBS: Diagnostic BioSystems : BD: Becton Dickinson Biosciences
